# Expression of fatty acyl-CoA ligase drives one-pot de novo synthesis of membrane-bound vesicles in a cell free transcription-translation system

**DOI:** 10.1101/2021.04.22.440977

**Authors:** Ahanjit Bhattacharya, Christy J. Cho, Roberto J. Brea, Neal K. Devaraj

## Abstract

Despite the central importance of lipid membranes in cellular organization, it is challenging to reconstitute their de novo formation from minimal chemical and biological elements. Here we describe a chemoenzymatic route to membrane-forming non-canonical phospholipids in which cysteine-modified lysolipids undergo spontaneous coupling with fatty acyl-CoA thioesters generated enzymatically by a fatty acyl-CoA ligase. Due to the high efficiency of the reaction, we were able to optimize phospholipid membrane formation in a cell-free transcription-translation (TX-TL) system. Combining DNA encoding for the fatty acyl-CoA ligase with suitable lipid precursors, enabled spontaneous one-pot de novo synthesis of membrane-bound vesicles. Non-canonical sphingolipid synthesis was also possible by using a cysteine-modified lysosphingomyelin as a precursor. When the sphingomyelin-interacting protein lysenin is co-expressed alongside the acyl CoA ligase, the in situ assembled membranes were spontaneously modified with protein. Our strategy of coupling gene expression with membrane lipid synthesis in a one-pot fashion could facilitate the generation of proteoliposomes and brings us closer to the bottom-up generation of synthetic cells using recombinant synthetic biology platforms.

## Introduction

There is a growing interest in the generation of compartmentalized structures to provide chassis for artificial cells.^1,2^ Lipids are arguably the most widely used building blocks for such compartments.^3–5^ Over the past decade, there has been a significant effort towards developing biomimetic lipid synthesis strategies.^6–8^ In living cells, phospholipids constitute the major class of membrane-forming lipids. Phospholipids are synthesized in part by membrane-bound acyltransferases, which couple various lysophospholipids with long-chain fatty acyl thioesters (Figure 1A).^9,10^ Integration of lipid synthesis with cell-free TX-TL has been a highly sought-after goal in synthetic cell development and membrane mimetic chemistry.^11^ In the past, there has been considerable effort to reconstitute bacterial phospholipid synthesis pathways in cell-free systems^12–14^. However, reconstitution of the natural phospholipid synthesis pathways in artificial cells is highly challenging for many reasons. The required membrane-bound protein enzymes can be difficult to express and fold in functional form. Also, the natural pathways consist of many biochemical components and steps which demand extensive optimization for reconstitution in cell-free conditions. These issues become exacerbated in simplified cell-free expression systems, which do not typically benefit from the machinery needed to ensure proper insertion and folding of transmembrane proteins. Furthermore, de novo vesicle formation is not possible since many of the required proteins need preexisting membranes for proper function. For these reasons, we became interested in exploring simpler strategies for membrane lipid synthesis that would utilize soluble biochemical elements and a minimal number of chemical steps. Simplified non-canonical lipid membrane generation would potentially facilitate future integration with bottom-up synthetic biology applications such as the creation of proteoliposomes and artificial cells.^6^

**Figure 1.**
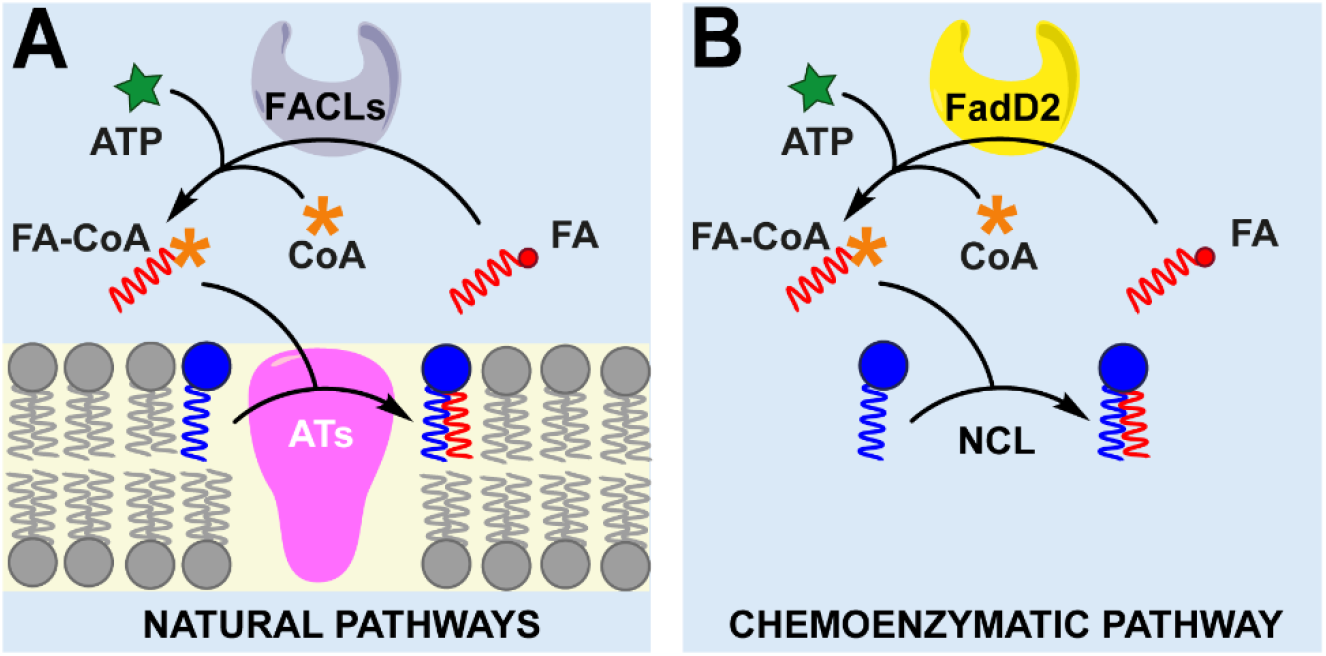
Phospholipid synthesis by ligation between single-chain amphiphiles. (A) Natural pathways. (B) Artificial pathway combining an enzymatic reaction with chemical ligation step. [AT: acyltransferases, FACL: fatty acyl CoA ligase, FA: fatty acid, FA-CoA: fatty acyl coenzyme A, NCL: native chemical ligation].

Here we describe a novel lipid synthesizing system by repurposing the activity of fatty acyl CoA ligases (FACLs). In particular, we utilized a mycobacterial FACL, FadD2,^15^ to catalyze the formation of fatty acyl CoAs (FA-CoA). FA-CoAs subsequently undergo chemoselective native chemical ligation (NCL) with cysteine-modified lysolipids to produce membrane-forming phospholipids (Figure 1B). By combining enzymatic and chemical steps, we preclude the need for a membrane-bound acyltransferase enzyme which is typically necessary to phospholipid synthesis. The high efficiency of the chemoenzymatic reaction enabled optimization of the reaction such that it was compatible in cell-free TX-TL systems. Analogues of both phosphatidylcholines and sphingomyelins can be generated. We demonstrate that FadD2 expression in a cell-free TX-TL system (PURE)^16^ can be couple to the spontaneous formation of membrane-bound vesicles. Finally, we show that the scope of our method can be expanded to the co-expression of both the acyl-CoA ligase and a membrane-interacting protein, such that protein modified vesicles can form in one-pot from the TX-TL system. Our results lay the groundwork for further elaboration of minimal expression systems capable of spontaneously forming functional membrane compartments.

## Results and discussion

### FadD2 drives the synthesis of cysteine-modified phospholipids

In a previous work, we demonstrated that cysteine-modified lysolipids can be coupled to amphiphilic fatty acyl thioesters by Native Chemical Ligation (NCL) under mild aqueous conditions to generate non-canonical phospholipids which subsequently self-assemble to form membrane-bound vesicles.^17^ NCL-driven vesicle formation has been applied to remodel artificial membranes^18^ and reconstitute functional transmembrane proteins.^19^ In biology, FACLs convert fatty acids into coenzyme A thioesters (FA-CoA) in two steps: (i) in the first step, the fatty acid is activated to the corresponding adenylate with ATP and Mg^2+^; (ii) in the second step, the enzyme-bound fatty acyl adenylate is converted to the corresponding FA-CoA and released from the active site.^20^ As they are soluble proteins, FACLs are convenient to express in functional form and purify. Also, there is an enormous diversity of FACLs in living organisms and many choices are available to meet specific substrate or kinetic requirements.^21^ We hypothesized that an enzymatically generated FA-CoA would spontaneously react with a cysteine-modified lysolipid by NCL (Figure 2A). We expressed and purified N-terminal His_6_-tagged mycobacterial FadD2 adapting a previously published procedure.^15^ To test the FACL activity, we incubated FadD2 with sodium dodecanoate (Na-DDA), MgCl_2_, ATP, CoA and tris-(2-carboxyethyl)phosphine (TCEP) at 37 °C and observed the formation of dodecanoyl-CoA (m/z = 948.26 for [M-H]^-^) by high performance liquid chromatography-mass spectrometry (HPLC-MS) (Figure 2B). When 1 equivalent of cysteine-modified lysolipid **L1** (see Figure S1a-b and Supporting Information for synthesis and characterization) was included in the reaction medium, we detected formation of the Cys-phospholipid **P1** (see Figure S1c-e and Supporting Information for full characterization). Differential scanning calorimetry (DSC) experiments indicate that **P1** forms fluid phase membranes under the experimental conditions (Figure S1f). If dodecanoic acid is added in molar excess of the lysolipid **L1**, the free thiol group on **P1** undergoes acylation to generate the three-tailed phospholipid **P1’**, as identified by mass spectrometry (Figure S2a). We also tested several unsaturated (C14:1, C16:1, C18:1) fatty acids in our chemoenzymatic setup and in all cases detected formation of the corresponding Cys-phospholipids by HPLC-MS (Table S1).

**Figure 2.**
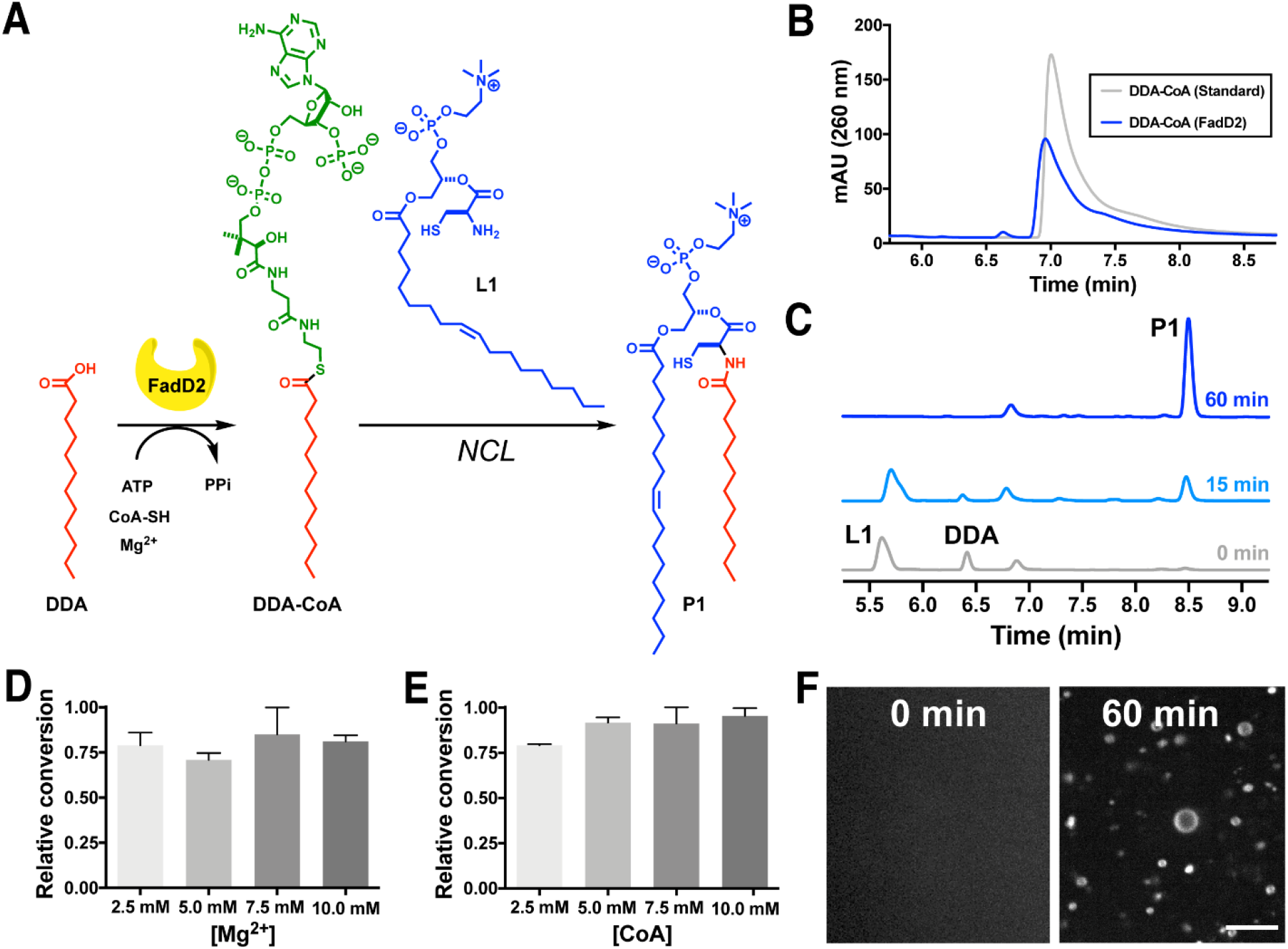
FadD2-driven phospholipid synthesis. (A) Reaction scheme corresponding to cysteine-modified phospholipid formation. (B) HPLC chromatogram (260 nm) corresponding to dodecanoyl-CoA generated by FadD2 (blue trace). For comparison, the chromatogram corresponding to a dodecanoyl-CoA standard is shown (gray trace). (C) HPLC-ELSD chromatograms at sequential time points showing progress of phospholipid **P1** formation. (D) Comparison of formation of phospholipid **P1** at various concentrations of Mg^2+^. (E) Comparison of formation of phospholipid **P1** at various concentrations of CoA. (F) Spinning disk confocal microscopy of *in situ* phospholipid membrane formation. Membranes were stained with 0.05 mol% Texas Red-DHPE. Scale bar: 10 μm.

Initially, we sought to determine the optimum conditions for FadD2-driven synthesis of phospholipids. We incubated lysolipid **L1**, Na-DDA, MgCl_2_, ATP, CoA, TCEP and FadD2 at various pH (7.0, 7.5, 8.0, 8.5) and followed the progress of the reactions by high performance liquid chromatography-mass spectrometry in conjunction with a evaporative light scattering detector (HPLC-ELSD-MS). Below pH 7.5, the conversion was low, likely due to poor solubility of the fatty acid and weak nucleophilicity of the thiol and amine groups on cysteine. An increase in the degree of conversion could be observed with an increase in pH. However, raising the pH too much led to decomposition of the phospholipid product due to ester hydrolysis at the *sn2* position. As such, we observed an optimum yield of the phospholipid product at pH range between 7.5-8.0. We followed the progress of the reaction consisting of 1 mM lysolipid **L1**, 2 mM TCEP, 9 mM MgCl_2_, 2.5 mM ATP, 1.25 mM CoA, 0.12 mg/mL FadD2, and 1 mM Na-DDA in 100 mM Na-HEPES at pH 8.0 at various time points by HPLC-ELSD-MS and observed complete conversion of the lysolipid **L1** within 1 h (Figure 2C). We tested the dependence of phospholipid synthesis on Mg^2+^ concentration and found that conversion does not significantly change over a Mg^2+^ concentration range of 2.5-10 mM (Figure 2D). Since CoA is regenerated over the course of the reaction, we also tested the dependence of phospholipid formation on the concentration of CoA. We found that above 50 mol% CoA (with respect to fatty acid), the extent of conversion is nearly unchanged (Figure 2e). Finally, we followed the progress of reaction using microscopy to test if the in situ formed phospholipids spontaneously self-assembled into vesicles. The reaction mixture was clear upon mixing all components and after 1 h, micron-sized vesicles were readily detected by fluorescence microscopy using a membrane staining dye (Figure 2F, Figure S3).

To test if alternative phospholipids can be formed, we synthesized a cysteine-modified lysosphingomyelin **L2** (see Figure S4a-b and Supporting Information for full characterization). When we incubated **L2** with 2 mM TCEP, 9 mM MgCl_2_, 2.5 mM ATP, 1.25 mM CoA, 0.12 mg/mL FadD2, and 1 mM Na-DDA in 100 mM Na-HEPES at pH 8.0 at 37 °C, we observed the efficient formation of the corresponding Cys-sphingolipid **P2** (Figure S5a). A small amount of the double acylation product **P2’** (Figure S2b) was also observed by HPLC-MS. Micron-sized vesicles were identified in the reaction mixture by microscopy (Figure S5b). To confirm **P2** alone can form vesicles, we carried out chemical synthesis of the compound (Figure S4c, Supporting Information for full synthetic details and characterization). We found from DSC experiments that **P2** exists in a fluid phase at room temperature (Figure S4d). Using optical microscopy and negative-staining transmission electron microscopy (TEM) we found that a thin film of purified **P2** forms membrane-bound vesicles when hydrated (Figure S4e-f).

### Optimization of lipid synthesis in PURE system

The PURE (protein synthesis using recombinant elements) system is a commercially available TX-TL system composed of recombinant proteins, tRNAs, ribosomes and small molecule factors.^16^ The well-defined nature of the PURE system makes it suitable for applications requiring the generation of components and building blocks for synthetic cells.^2^ The PURE system is compatible with various pre-formed lipidic structures such as liposomes,^22^ nanodiscs,^23^ sponge droplets,^24^ and supported lipid bilayers.^25^ However, such pre-formed structures are passive components and are unlikely to directly interfere with the chemistry of the PURE system. A more challenging goal is to integrate active lipid synthesis pathways into the PURE system. Lipid synthesis could potentially perturb the biochemistry of the PURE system and vice versa. Therefore, we sought to determine whether the FACL-driven lipid synthesis can be combined with the PURE system to carry out simultaneous protein expression and membrane formation in a ‘one-pot’ fashion (Figure 3A).

**Figure 3.**
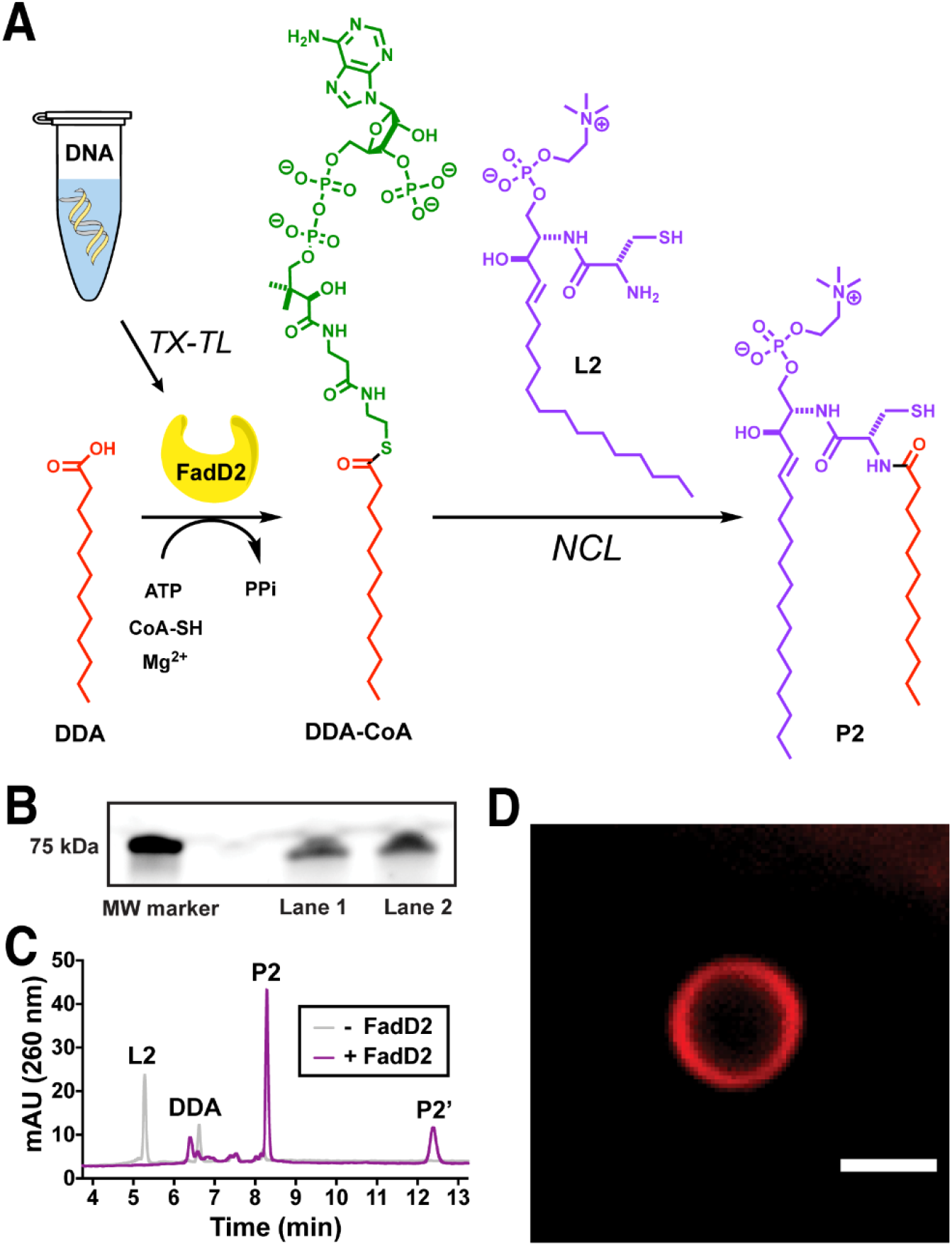
One-pot synthesis of membrane-forming lipids in PURE system mediated by FadD2. (A) Reaction scheme corresponding to cysteine-modified sphingomyelin formation driven by FadD2 expressed in PURE system. (B) SDS-PAGE (in-gel fluorescence) showing expression of FadD2 labeled with BODIPY-lysine. Lane 1: no lipid precursors, Lane 2: lipid precursors added. (C) HPLC-ELSD chromatograms showing the formation of **P2** under one-pot condition. (D) Spinning disk confocal microscopy of vesicles formed from **P2** under one-pot condition in PURE system. Scale bar represents 2 μm.

As an initial test, we simply combined the aforementioned lipid synthesis precursors and FACL-encoding DNA with the PURExpress kit (New England Biolabs) components. We added the FadD2 linear DNA template, lysolipid **L1** (0.48 mM), Na-DDA (0.48 mM), ATP (2.38 mM), CoA (0.60 mM), and TCEP (2.38 mM) to the PURE system components. Interestingly, we did not observe any protein expression nor synthesis of artificial phospholipid. In an effort to make TX-TL and lipid synthesis mutually compatible, we first examined which components in the lipid synthesis reaction were responsible for the deleterious effect on TX-TL. To do this, we followed fluorometrically the expression of sfGFP in the presence of each of the components required for chemoenzymatic lipid synthesis (Figure S6). We determined that TX-TL is completely inhibited when extra ATP is added (Figure S6). Inhibition of TX-TL likely happens because the excess ATP sequesters the critically important Mg^2+^ present in the PURE system.^2^ We found that this issue could be solved by adding an equivalent amount of Mg^2+^. We also found that lysolipids such as **L1**, which are modified with an α-amino acid are hydrolytically unstable at the *sn2* ester linkage in the undiluted PURE reaction, likely due to the presence of high concentration of nucleophiles, such as amino acids. So, we chose to work with the lysolipid **L2**, which is hydrolytically stable due to presence of amide linkage.

Having optimized the conditions for sfGFP expression, we then attempted the expression of FadD2 by adding a premix containing optimal concentrations of ATP (2.09 mM), CoA (0.698 mM), Mg^2+^ (2.56 mM), TCEP (2.33 mM), lysolipid **L2** (0.93 mM) and Na-DDA (0.93 mM). To test the expression of FadD2, we supplemented the reaction with a fluorescently modified lysyl-tRNA (FluoroTect^TM^ Green_Lys_, Promega) which incorporates a fluorescent probe into expressed proteins during translation. Combining SDS-PAGE with in-gel fluorescence detection, we confirmed that the expression of FadD2 took place in the presence of the **P2** lipid precursors (Figure 3B). We then tested the one-pot expression of FadD2 and chemoenzymatic lipid formation and observed formation of lipid **P2** by HPLC-ELSD-MS in high yield, with no remaining **L2** detectable (Figure 3C). A small amount of the diacylated product **P2’** was also detected. We observed micron-sized vesicles in the same reaction mixture using spinning disk confocal microscopy (Figure 3D, Figure S7a). Smaller vesicles on the order of 100 nm were observed by negative staining TEM (Figure S7b). In comparison, when the FadD2 DNA is omitted from the PURE reaction, no lipid synthesis occurred, and no vesicles were detected by TEM (Figure S7c) or optical microscopy (Figure S7d).

### Co-expression of membrane-interacting proteins

Finally, we tested if a membrane-interacting protein can be co-expressed and FadD2-driven lipid membrane formation can be achieved under the optimized one-pot condition. Since we are generating a modified-sphingomyelin (**P2**), we chose sfGFP-NT-lysenin (NT: non-toxic) as a model protein for this experiment. NT-lysenin is a truncated form of the sphingomyelin-interacting protein lysenin that has been shown to selectively bind to sphingomyelins over other sphingolipid species.^26–28^ We first carried out the expression of sfGFP-NT-lysenin in the presence of pre-formed vesicles of purified **P2**. We observed that the protein is indeed localized to the membranes, suggesting that NT-lysenin can bind to the sphingomyelin **P2** even with the cysteine-modification (Figure 4A). When sfGFP-NT-lysenin was expressed in presence of dioleoylphosphatidylcholine (DOPC) vesicles, no protein localization to the membranes was observed suggesting that the expressed sfGFP-NT-lysenin does not have any non-specific interactions with lipid membranes (Figure 4B). Having obtained these results, we carried out TX-TL reaction in the presence of FadD2 and sfGFP-NT-lysenin DNA templates and lipid precursors (Figure 4C). We observed the formation of the sphingolipid **P2** by HPLC-MS (Figure S8a). To confirm that both proteins were expressed, we supplemented the reaction with a fluorescently modified lysyl-tRNA (FluoroTect^TM^ Green_Lys_). Using SDS-PAGE and in-gel fluorescence detection, we confirmed the expression of both proteins in the presence of the **P2** lipid precursors (Figure 4D). When the one-pot condition was tested with co-expression of FadD2 and sfGFP-NT-lysenin, we observed vesicles with the fluorescent protein bound to the membranes (Figure 4E, Figure S8b).

**Figure 4.**
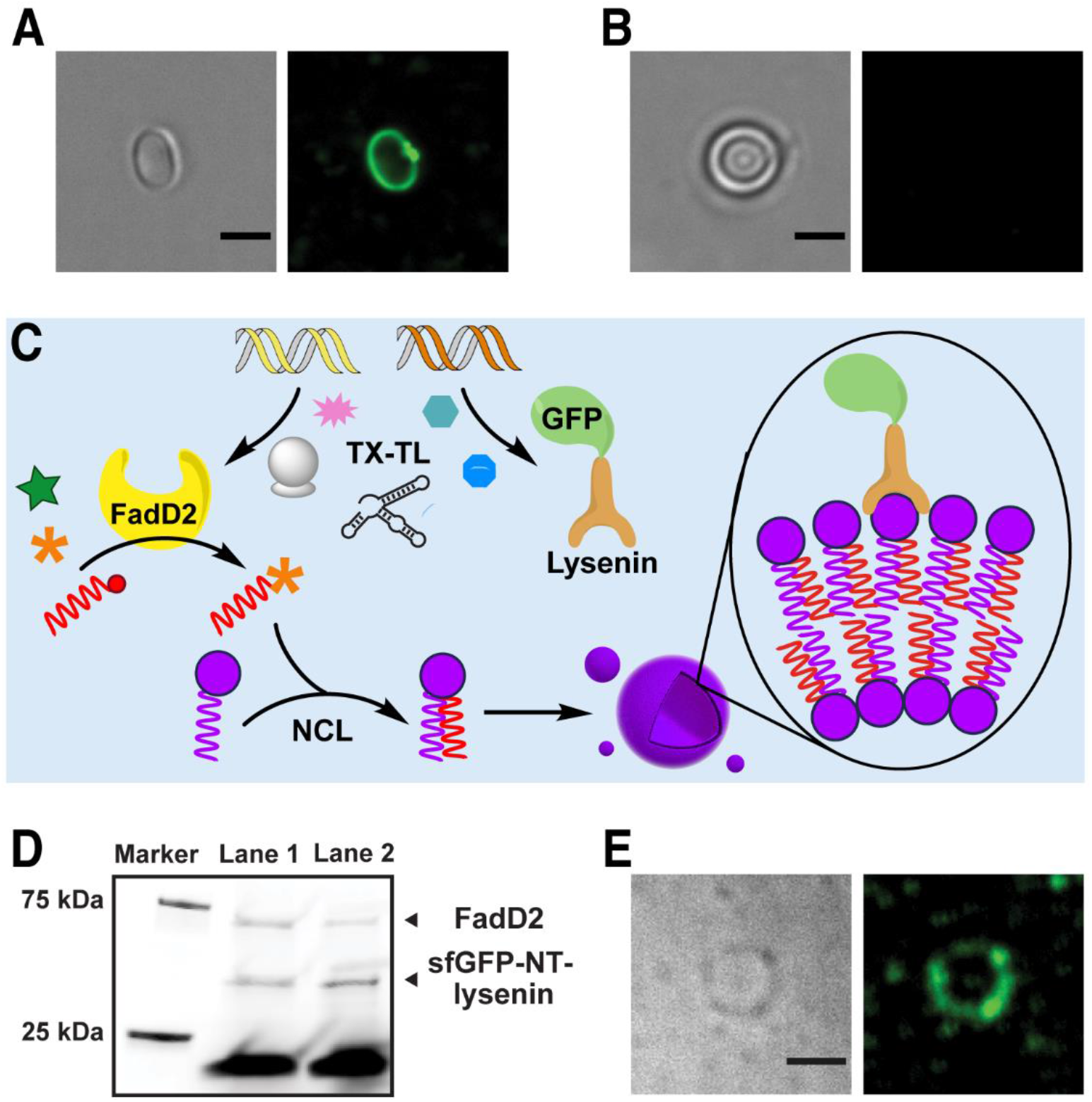
Localization of a membrane-binding protein to in situ formed lipid membranes under one-pot condition. (A) Expression of sfGFP-NT-lysenin in presence of sphingomyelin **P2** vesicles. (B) Expression of sfGFP-NT-lysenin in presence of DOPC vesicles. (C) Schematic representation outlining the concept of one-pot expression of the fatty acid activating enzyme FadD2 and a membrane-binding protein sfGFP-NT-lysenin and subsequent localization of the latter to the in situ formed membranes. (D) SDS-PAGE (in-gel fluorescence) showing expression of FadD2 and sfGFP-NT-lysenin labeled with BODIPY-lysine. Lane 1: no lipid precursors, Lane 2: lipid precursors added. (E) A sphingomyelin **P2** vesicle generated under one-pot condition and localizing sfGFP-NT-lysenin to the membranes. All scale bars represent 3 μm.

## Conclusion

In summary, we have described a chemoenzymatic method to generate cell-like membrane compartments in a cell-free TX-TL system. Recently we reported a minimal biochemical route to membrane-forming phospholipids by repurposing the activity of a soluble mycobacterial fatty acyl adenylate ligase (FAAL) FadD10.^29^ There, we synthesized membrane-forming diacylphospholipids through chemoselective coupling of enzymatically generated fatty acyl adenylates with amine-modified lysolipids. We expressed FadD10 in the PURE system, and subsequently added lipid precursors to drive phospholipid synthesis. However, FadD10-driven phospholipid synthesis could not be efficiently implemented in a one-pot fashion. In this work, we overcame that roadblock by working with FACLs. The high chemoselectivity and fast kinetics of coupling between enzymatically generated FA-CoAs and cysteine-modified lysolipids ensured that undesired reactions with potential competing groups are minimized. To our knowledge, chemoenzymatic de novo generation of lipid membranes simultaneously with protein synthesis has not been previously demonstrated in cell-free TX-TL systems. Thus, our results will further our understanding of how cell-free systems function, particularly in the presence of multiple exogenous molecules. The facile implementation of a lipid synthesis strategy with commercially available TX-TL systems such as the PURE system will enable the modular integration of lipid synthesis pathways into genetic circuits. We further demonstrated that membrane-interacting proteins can be co-expressed with the FACL and localized onto the in situ synthesized lipid membranes. In a similar manner, many functional proteins can be localized to the membrane surface based on well-defined affinity interactions. We envision that, if the efficiency of TX-TL systems undergoes further improvement in the future, many more proteins may be expressed simultaneously, and complex biochemical pathways may be reconstituted. Finally, simplified biochemical routes to membrane-forming lipids and proteoliposomes can provide us hints at how early cellular membrane synthesizing machinery evolved.

## Materials and Methods

### General Considerations

All reagents obtained from commercial suppliers were used without further purification unless otherwise noted. 1-oleoyl-2-hydroxy-*sn*-glycero-3-phosphocholine (Lyso C_18:1_ PC-OH) and dioleoylphosphatidylcholine (DOPC) were obtained from Avanti^®^ Polar Lipids. Sphingosylphosphorylcholine [Lyso-sphingomyelin (d18:1); Lyso d_18:1_ SM-NH2], coenzyme A, and tris(2-carboxyethyl) phosphine (TCEP) were obtained from Cayman Chemicals. *N*-Boc-L-Cys(Trt)-OH, *N*-Fmoc-L-Cys(Trt)-OH, 2,4,6-trichlorobenzoyl chloride (TCBC), 4-dimethylaminopyridine (DMAP), O-(7-azabenzotriazol-1-yl)-1,1,3,3-tetramethyl-uronium hexafluorophosphate (HATU), adenosine triphosphate (ATP), dichloromethane (CH_2_Cl_2_), *N,N*-dimethylformamide (DMF), *N,N*-diisopropylethylamine (DIEA), trifluoroacetic acid (TFA), triethylsilane (TES), 4-methylpiperidine, dodecanoic acid, and lysozyme (chicken egg white) were obtained from Sigma-Aldrich. Texas Red^®^ 1,2-dihexadecanoyl-*sn*-glycero-3-phosphoethanolamine, triethylammonium salt (Texas Red^®^ DHPE) and Nile Red were obtained from Life Technologies. Deuterated chloroform (CDCl_3_) and methanol (CD_3_OD) were obtained from Cambridge Isotope Laboratories. Proton nuclear magnetic resonance (^1^H NMR) spectra were recorded on a Varian VX-500 MHz spectrometer, and were referenced relative to residual proton resonances in CDCl_3_ (at δ7.24 ppm) or CD_3_OD (at δ4.87 or δ3.31 ppm). Chemical shifts were reported in parts per million (ppm, δ) relative to tetramethylsilane (δ0.00). ^1^H NMR splitting patterns are assigned as singlet (s), doublet (d), triplet (t), quartet (q) or pentuplet (p). All first-order splitting patterns were designated on the basis of the appearance of the multiplet. Splitting patterns that could not be readily interpreted are designated as multiplet (m) or broad (br). Carbon nuclear magnetic resonance (^13^C NMR) spectra were recorded on a Varian VX-500 MHz spectrometer, and were referenced relative to residual proton resonances in CDCl_3_ (at δ77.23 ppm) or CD_3_OD (at δ49.15 ppm). Electrospray Ionization-Time of Flight (ESI-TOF) spectra were obtained on an Agilent 6230 Accurate-Mass TOF-MS mass spectrometer. Spinning-disk confocal microscopy images were acquired on a Yokagawa spinning-disk system (Yokagawa, Japan) built around an Axio Observer Z1 motorized inverted microscope (Carl Zeiss Microscopy GmbH, Germany) with a 63x, 1.40 NA oil immersion objective to an Orca Flash 4.0 CMOS camera (Hamamatsu) using ZEN imaging software (Carl Zeiss Microscopy GmbH, Germany). NanoDrop 2000C spectrophotometer was used for UV/Vis measurements. Fluorescence measurements were carried out on a Tecan Infinite F200 plate reader.

### HPLC methods

HPLC experiments were performed on an Agilent 1260 Infinity LC System. All analytical runs were performed using an Eclipse Plus C8 analytical column with an evaporative light scattering detector (ELSD) at a flow rate of 1.0 mL/min. ELSD enabled sensitive detection of lipids. HPLC purification was carried out on Zorbax SB-C18 semipreparative column at a flow rate of 4.0 mL/min. Various ratios of *Phase A* (H_2_O with 0.1% v/v formic acid) and *Phase B* (MeOH with 0.1% v/v formic acid) were used as mobile phase. Retention times (t_R_ were verified by mass spectrometry.

### Critical micelle concentration (cmc) determination

The cmc’s of the lysolipids **L1** and **L2** were determined using a method based on the solvatochromic fluorescent dye Laurdan.^30^ Laurdan shows a sharp change in the value of generalized polarization (GP) at or near the cmc value of an amphiphile. Initially, a 10 mM solution of the lysolipid was diluted to various concentrations over the range 1-500 μM. 2.5 mM TCEP was included in each solution to ensure that the thiol groups stay reduced. The final concentration of Laurdan was 3 μM in each dilution. Then, the samples were transferred to a 384 well plate and analyzed on a Tecan Infinite Plate Reader at 37 °C. The samples were excited at 364 nm and emission spectra acquired over 430-500 nm. GP was calculated as follows:

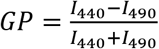

where *I*_440_ and *I*_490_ stands for the fluorescence intensities at 440 nm and 490 nm, respectively. The values of GP were plotted against the lipid concentrations for each dilution.

### Differential scanning calorimetry (DSC)

DSC experiments were carried out on a Microcal VP-Capillary instrument to study the gel-to-fluid phase transition behavior of the cysteine-modified lipids **P1** and **P2**. Aqueous dispersions of the lipids were prepared at concentrations of 0.25 mM in Milli-Q H_2_O containing 1 mM TCEP. 450 μL of the dispersion was taken for individual scans. The scan rate was 30 °C/h and *gain* was set to “high”. Manufacturer provided Microcal Origin Software was used to process the data.

### Transmission electron microscopy (TEM)

4.8 μL of a test sample was added to formvar-coated Cu grid surface and allowed to sit for ~1 min. Afterwards the excess solution was blotted into a filter paper. The grid was washed twice with 4.8 μL of 100 mM HEPES-K pH 7.6 (for PURE reaction) or ultrapure H_2_O (for **P1** and **P2** vesicles). After this 4.8 μL drops of 2% uranyl acetate were added and quickly blotted three times. The stain was left for ~30 s in the fourth time following which excess stain was blotted. The grids were dried in air. Finally, the grids were imaged by TEM on an FEI Tecnai Spirit G2 BioTWIN microscope equipped with an Eagle 4k (16 megapixel) camera.

### Expression and purification of FadD2^15^

The *pET28b-fadD2* (*Rv0270*) and pGro7 (chaperone, Takara) plasmids were co-transformed into BL21(DE3) competent *E. coli* cells (New England Biolabs) and colonies were generated on a Luria-Bertani (LB) agar plate containing kanamycin and chloramphenicol. A single colony was picked and grown overnight at 37 °C in LB broth containing kanamycin (0.05 mg/mL) and chloramphenicol (0.025 mg/mL). Afterward, 10 mL of the overnight culture was used to inoculate 1 L of freshly autoclaved LB medium containing kanamycin (0.05 mg/mL), chloramphenicol (0.025 mg/mL), and L-arabinose (0.5 mg/mL). The culture was grown at 37 °C in a shaker-incubator till the OD600 reached about 0.7. Over-expression of FadD2 was induced by addition of 1 mM isopropyl 1-thio-D-galactopyranoside (IPTG) after bringing the culture to room temperature. The cells were then grown for 20 h at 18 °C, after which the culture was centrifuged at 6,000 rcf for 20 min at 4 °C. The pellet was resuspended by vortexing in 12 mL of lysis buffer containing 50 mM Na-phosphate pH 8.0, 10 mM imidazole pH 8.0, 0.5 M NaCl, 2 mM β-mercaptoethanol, 1 mg/mL lysozyme and PMSF (1 mM) and kept on ice for 30 min. After lysing the cells using an ultrasonicator probe, debris were removed by centrifuging (10,000 rcf, 25 min, 4 °C). The supernatant was incubated in a gravity column with Ni^2+^-nitrilotriacetate (Ni-NTA) agarose resin on a shaker table for 1.5 h at 4 °C. Next, the flow-through was discarded and the resin was washed with 25 mM, 35 mM, 40 mM, and 50 mM imidazole solutions (each containing 50 mM Na-phosphate pH 8.0 and 0.1 M NaCl). Finally, His_6_-tagged FadD2 was eluted with 250 mM imidazole (containing 50 mM Na-phosphate pH 8.0 and 0.1 M NaCl) and collected in 1 mL fractions. The pure fractions were pooled, washed (with 50 mM Na-phosphate pH 8.0 and 0.1 M NaCl) and concentrated using 30 kDa and 50 kDa MWCO centrifuge filters at 4 °C. The enzyme (2.4 mg/mL) was stored at 4 °C.

### Assays on FadD2-mediated phospholipid synthesis

A typical phospholipid synthesis reaction consisted of 1 mM lysolipid (**L1** or **L2**), 2 mM TCEP, 9 mM MgCl_2_, 2.5 mM ATP, 1.25 mM CoA, 0.12 mg/mL FadD2, and 1 mM Na-dodecanoate in 100 mM Na-HEPES pH 8.0 and it was incubated at 37 °C. For LC-MS analysis, the reaction was quenched with 90 μL MeOH and 0.5 μL 50% β-mercaptoethanol and insoluble materials were removed by centrifugation. The supernatant was carefully collected and injected into HPLC. In the experiments where the effect of variable concentrations of Mg^2+^ and CoA were tested, the reactions were quenched after 30 min and analyzed by LC-MS. For microscopic observations, 0.05 mol% Texas Red-DHPE – a lipophilic dye was added in the reaction mixture.

### Synthesis of membrane-forming lipids in PURE system

*Preparation of linear DNA templates*. Linear DNA templates were used for all transcription-translation (TX-TL) reactions. Briefly, the templates were prepared by polymerase chain reaction (PCR) amplification from corresponding plasmid constructs using Q5 DNA Polymerase Master Mix (NEB). The PCR reaction was worked up using a Qiagen PCR purification kit. The DNA was eluted in MilliQ H_2_O and stored at −20 °C. The size and purity of the linear template were verified by agarose gel electrophoresis.

#### Preparation of premix of lipid precursors

As the PURE system reaction can accommodate only a small volume (~20%) excess of the manufacturer-guided reaction volume, we added the extra components from concentrated ‘premix’ solutions for reliable pipetting. A typical premix of lipid precursors (10 μL volume) was prepared by adding the following components: lysolipid **L2** (20 mM) – 2 μL (4 mM), TCEP (100 mM) – 1 μL (10 mM), ATP (100 mM) – 0.9 μL (9 mM), CoA (10 mM) – 3 μL (3 mM), MgCl_2_ (100 mM) – 1.1 μL (11 mM), Na-DDA (20 mM) – 2 μL (4 mM). All individual stock solutions were prepared in nuclease free H_2_O. The premix was stored at −20 °C and used within 2 days.

#### HPLC analysis of one-pot reaction and microscopic observation

A PURE reaction was set up by adding the following: Solution A (4 μL), Solution B (3 μL), murine RNase inhibitor (0.25 μL, 40 U/μL), FadD2 linear DNA (12.8 nM final), lipid precursors premix (final concentrations: lysolipid **L2** – 0.930 mM, TCEP – 2.33 mM, ATP – 2.09 mM, CoA – 0.698 mM, MgCl_2_ – 2.56 mM, Na-DDA – 0.930 mM. The final volume of the reaction was 10.75 μL. For HPLC detection, the reaction was quenched with 90 μL MeOH and 0.5 μL 50% β-mercaptoethanol and the white precipitate generated was removed by centrifugation. The clear supernatant was injected into HPLC for analysis. For microscopic observation, the reaction mixture was added onto a dried film of Nile Red and gently tapped. A small volume was placed on a glass slide, secured with a cover slip and imaged by spinning disk confocal microscopy.

#### Fluorescent detection of protein expression

A PURE reaction was set up by adding the following: Solution A (4 μL), Solution B (3 μL), murine RNase inhibitor (0.25 μL, 40 U/μL), FluorTect^TM^ Green_Lys_ (0.2 μL), FadD2 linear DNA (10.2 nM final), lipid precursors premix (final concentrations: lysolipid **L2** – 0.375 mM, TCEP – 1.03 mM, ATP – 0.845 mM, CoA – 0.117 mM, MgCl_2_ – 0.939 mM, Na-DDA – 0.375 mM. In a control reaction, no lipid precursors were added. The final volumes of the reactions were 10.65 μL. The reactions were kept incubated at 37 °C for 2 h. The entire reaction mixtures were loaded on a pre-cast 4-20% polyacrylamide gel and separated electrophoretically. The gel was imaged on a Typhoon^TM^ FLA 9500 scanner (473 nm laser, Cy2 channel).

### Co-expression of FadD2 and sfGFP-NT-lysenin in PURE system

*HPLC detection of lipid synthesis and microscopic observation*. A PURE reaction was set up by adding the following: Solution A (4 μL), Solution B (3 μL), murine RNase inhibitor (0.25 μL, 40 U/μL), FadD2 linear DNA (12.8 nM final), sfGFP-NT-lysenin linear DNA (2.4 nM final), lipid precursors premix (final concentrations: lysolipid **L2** – 0.881 mM, TCEP – 2.20 mM, ATP – 1.98 mM, CoA – 0.661 mM, MgCl_2_ – 2.42 mM, Na-DDA – 0.881 mM. The final volumes of the reactions were 11.35 μL. For HPLC detection, the reaction was quenched with 90 μL MeOH and 0.5 μL 50% β-mercaptoethanol. The insoluble aggregates are precipitated by centrifugation and the supernatant was analyzed by HPLC-MS. For microscopic observation, a small volume of the reaction was taken on a glass slide, secured with a cover slip and imaged by spinning disk confocal microscopy. *Fluorescent detection of protein expression*. A PURE reaction was set up by adding the following: Solution A (4 μL), Solution B (3 μL), murine RNase inhibitor (0.25 μL, 40 U/μL), FluorTect^TM^ Green_Lys_ (0.2 μL), FadD2 linear DNA (10.2 nM final), sfGFP-NT-lysenin linear DNA (7.7 nM final), lipid precursors premix (final concentrations: lysolipid **L2** – 0.375 mM, TCEP – 1.03 mM, ATP – 0.845 mM, CoA – 0.117 mM, MgCl_2_ – 0.939 mM, Na-DDA – 0.375 mM. In a control reaction, no lipid precursors were added. The final volumes of the reactions were 10.65 μL. The reactions were kept incubated at 37 °C for 2 h. The entire reaction mixtures were loaded on a pre-cast 4-20% polyacrylamide gel and separated electrophoretically. The gel was imaged on a Typhoon^TM^ FLA 9500 scanner (473 nm laser, Cy2 channel).

## Acknowledgements

This material is based upon work supported by the Department of Defense (Army Research Office) through the Multidisciplinary University Research Initiative (MURI) under Award No. W911NF-13-1-0383. We thank Anthony D Baughn (University of Minnesota) and Cuiwen He (University of California Los Angeles) for generously providing the plasmids for FadD2 and NT-lysenin respectively. We thank Gabriel Ozorowski (Ward Lab, The Scripps Research Institute) for help with DSC, Henrike Niederholtmeyer and Cynthia Chaggan for constructing the sfGFP-NT-lysenin plasmid.

## Notes

### Competing Interest Statement

The authors have declared no competing interest.

