## Supporting Information for "Expression of fatty acyl-CoA ligase drives one-pot de novo synthesis of membrane-bound vesicles in a cell free transcription-translation system"

Synthetic procedures……………………………………………………………………………..S2

Supplementary Tables…………………………………………………………………………….S6

Supplementary Figures…………………………………………………………………………...S7

Supplementary References……………………………………………………………………..S13

NMR Spectra……………………………………………………………………………….......S14

#### Synthetic procedures

Synthesis of cysteine-modified lysolipids

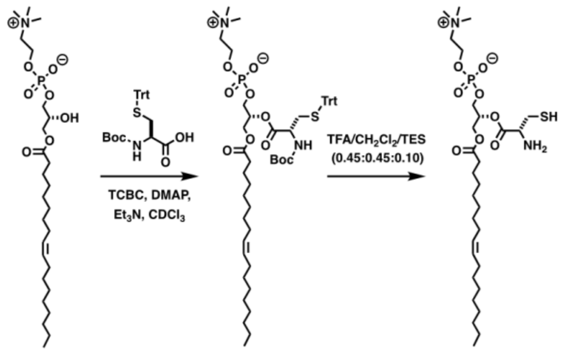


Lyso $C_{18:1}$ PC-OH    L1.1    L1

**1-oleoyl-2-[*N*-Boc-L-Cys(Trt)]-*sn*-glycero-3-phosphocholine (L1.1).**[1,2] A solution of 1-oleoyl-2-hydroxy-*sn*-glycero-3-phosphocholine (**Lyso $C_{18:1}$ PC-OH**, 50.0 mg, 95.8 µmol), *N*-Boc-L-Cys(Trt)-OH (111.1 mg, 239.6 µmol), DMAP (70.2 mg, 574.8 µmol) and $Et_3N$ (46.7 µL, 335.3 µmol) in $CDCl_3$ (3.8 mL) was stirred at rt for 10 min. Then, TCBC (97.3 µL, 622.7 µmol) was added. After 12 h stirring at rt, $H_2O$ (250 µL) was added to quench the acyl chloride, and the solvent was removed under reduced pressure to give a pale yellow solid. The corresponding residue was dissolved in MeOH (1 mL), filtered using a 0.2 µm syringe-driven filter, and the crude solution was purified by HPLC, affording 79.1 mg of lysophospholipid **L1.1** as a white solid [84%, $t_R$ = 7.8 min (100% *Phase B*, 15.5 min)]. $^1H$ NMR ($CDCl_3$, 500.13 MHz, δ): 7.38 (d, *J* = 7.5 Hz, 6H, 6 × $CH_{Ar}$), 7.33-7.26 (m, 6H, 6 × $CH_{Ar}$), 7.25-7.19 (m, 3H, 3 × $CH_{Ar}$), 5.40-5.29 (m, 2H, 2 × CH), 5.27-5.13 (m, 1H, 1 × CH), 5.07 (d, *J* = 9.0 Hz, 1H, 1 × NH), 4.41-3.88 (m, 7H, 3 × $CH_2$ + 1 × CH), 3.77-3.57 (m, 2H, 1 × $CH_2$), 3.25 (s, 9H, 3 × $CH_3$), 2.74-2.48 (m, 2H, 1 × $CH_2$), 2.31-2.09 (m, 2H, 1 × $CH_2$), 2.07-1.93 (m, 4H, 2 × $CH_2$), 1.61-1.43 (m, 2H, 1 × $CH_2$), 1.42 (s, 9H, 3 × $CH_3$), 1.31-1.17 (m, 20H, 10 × $CH_2$), 0.88 (t, *J* = 7.0 Hz, 3H, 1 × $CH_3$). $^{13}C$ NMR ($CDCl_3$, 125.77 MHz, δ): 173.5, 170.4, 163.9, 155.3, 144.4, 130.1, 129.9, 129.7, 129.6, 128.2, 128.2, 127.1, 80.1, 72.3, 67.2, 66.5, 63.8, 62.7, 59.4, 54.6, 52.7, 34.1, 34.1, 34.0, 32.0, 29.9, 29.9, 29.7, 29.5, 29.5, 29.4, 29.4, 29.3, 29.2, 29.2, 28.5, 28.5, 27.4, 27.3, 24.9, 24.8, 22.8, 14.3. MS (ESI-TOF) [m/z (%)]: 989 ($[M+Na]^+$, 20), 967 ($[MH]^+$, 100). HRMS (ESI-TOF) calculated for $C_{53}H_{80}N_2O_{10}PS$ ($[MH]^+$) 967.5266, found 967.5269.

**1-oleoyl-2-(L-Cys)-*sn*-glycero-3-phosphocholine (L1).**[1,2] A solution of 1-oleoyl-2-[N-Boc-L-Cys(Trt)]-*sn*-glycero-3-phosphocholine (**L1.1**, 10.0 mg, 10.3 µmol) in 1 mL of TFA/$CH_2Cl_2$/TES (0.45:0.45:0.10) was stirred at rt for 30 min. After removal of the solvent, the residue was dried under high vacuum for 3 h. Then, the corresponding residue was dissolved in MeOH (500 µL), filtered using a 0.2 µm syringe-driven filter, and the crude solution was purified by HPLC, affording 5.1 mg of the lysophospholipid **L1** as a colorless foam [79%, $t_R$ = 8.6 min (50% *Phase A* in *Phase B*, 5 min, and then 5% *Phase A* in *Phase B*, 10 min)]. $^1H$ NMR ($CD_3OD$, 500.13 MHz, δ): 5.46-5.26 (m, 3H, 3 × CH), 4.49-4.35 (m, 1H, 1 × CH), 4.34-4.19 (m, 3H, 1.5 × $CH_2$), 4.16-3.99 (m, 3H, 1.5× $CH_2$), 3.72-3.59 (m, 2H, 1 × $CH_2$), 3.29-3.26 (m, 1H, 0.5× $CH_2$), 3.23 (s, 9H, 3 × $CH_3$), 3.19-3.00 (m, 1H, 0.5× $CH_2$), 2.35 (t, *J* = 6.7 Hz, 2H, 1 × $CH_2$), 2.09-1.90 (m, 4H, 2 × $CH_2$), 1.69-1.53 (m, 2H, 1 × $CH_2$), 1.41-1.22 (m, 20H, 10 × $CH_2$), 0.90 (t, *J* = 6.7 Hz, 3H, 1 × $CH_3$). $^{13}C$ NMR ($CD_3OD$, 125.77 MHz, δ): 174.9, 172.3, 131.6, 130.8, 73.7, 67.4, 64.9, 63.4, 63.2, 60.6, 54.6, 34.9, 33.7, 33.1, 30.8, 30.8, 30.7, 30.6, 30.5, 30.4, 30.3, 30.3, 30.2, 28.1, 26.0, 23.8, 14.5. MS (ESI-TOF) [m/z (%)]: 625 ($[MH]^+$, 100). HRMS (ESI-TOF) calculated for $C_{29}H_{58}N_2O_8PS$ ($[MH]^+$) 625.3651, found 625.3647.

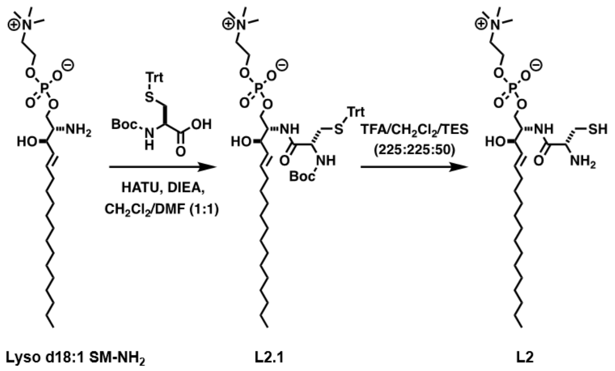

**Lyso-sphingomyelin (d18:1)-[*N*-Boc-L-Cys(Trt)] (L2.1).** A solution of *N*-Boc-L-Cys(Trt)-OH (6.0 mg, 12.9 µmol) in $CH_2Cl_2$/DMF (1:1) (250 µL) was stirred at 0 ºC for 10 min, and then HATU (5.4 mg, 14.2 µmol) and DIEA (9.0 µL, 51.7 µmol) were successively added. After 10 min stirring at 0 ºC, lyso-sphingomyelin (d18:1) (**Lyso $d_{18:1}$ SM-$NH_2$**, 6.0 mg, 12.9 µmol) was added. After 1 h stirring at rt, the mixture was concentrated under reduced pressure. The corresponding residue was dissolved in MeOH (500 µL), filtered using a 0.2 µm syringe-driven filter, and the crude solution was purified by HPLC, affording 9.7 mg of **L2.1** as a colorless film [83%, $t_R$ = 13.5 min (5% *Phase A* in *Phase B*, 15.5 min)]. $^1$H NMR ($CD_3OD$, 500.13 MHz, δ): 7.44-7.36 (m, 6H, 6 × $CH_{Ar}$), 7.35-7.28 (m, 6H, 6 × $CH_{Ar}$), 7.27-7.21 (m, 3H, 3 × $CH_{Ar}$), 5.69-5.55 (m, 1H, 1 × CH), 5.47-5.31 (m, 1H, 1 × CH), 4.34-4.17 (m, 2H, 1 × $CH_2$), 4.13-4.04 (m, 1H, 1 × CH), 4.03-3.96 (m, 2H, 1 × $CH_2$), 3.95-3.88 (m, 1H, 1 × CH), 3.87-3.81 (m, 1H, 1 × CH), 3.65-3.53 (m, 2H, 1 × $CH_2$), 3.19 (s, 9H, 3 × $CH_3$), 2.55-2.47 (m, 1H, 0.5 × $CH_2$), 2.46-2.39 (m, 1H, 0.5 × $CH_2$), 1.90-1.77 (m, 2H, 1 × $CH_2$), 1.45 (s, 9H, 3 × $CH_3$), 1.51-1.37 (m, 22H, 11 × $CH_2$), 0.89 (t, *J* = 7.0 Hz, 3H, 1 × $CH_3$). $^{13}$C NMR ($CD_3OD$, 125.77 MHz, δ): 172.5, 157.3, 146.0, 134.9, 130.8, 130.7, 129.1, 128.0, 80.7, 72.1, 68.0, 67.5, 65.5, 60.4, 55.4 and 55.2, 54.7, 36.0, 33.4, 33.1, 30.9, 30.8, 30.8, 30.7, 30.5, 30.4, 30.3, 28.8, 23.8, 14.5. MS (ESI-TOF) [m/z (%)]: 932 ([M+Na]$^+$, 16), 910 ([MH]$^+$, 100).

**Lyso-Sphingomyelin (d18:1)-[L-Cys] (L2).** A solution of lyso-sphingomyelin (d18:1)-[*N*-Boc-L-Cys(Trt)] (**L2.1**, 4.5 mg, 4.9 µmol) in 500 µL of TFA/$CH_2Cl_2$/TES (225:225:50) was stirred at rt for 30 min. After removal of the solvent, the residue was dried under high vacuum for 3 h. Then, the corresponding residue was dissolved in MeOH (250 µL), filtered using a 0.2 µm syringe-driven filter, and the crude solution was purified by HPLC, affording 2.2 mg of the lysosphingomyelin **L2** as a colorless film [78%, $t_R$ = 8.3 min (50-5% *Phase A* in *Phase B*, 5 min, and then 5% *Phase A* in *Phase B*, 10 min)]. $^1$H NMR ($CD_3OD$, 500.13 MHz, δ): 5.85-5.70 (m, 1H, 1 × CH), 5.56-5.43 (m, 1H, 1 × CH), 4.35-4.22 (m, 2H, 1 × $CH_2$), 4.15-4.01 (m, 3H, 1 × CH + 1 × $CH_2$), 3.98-3.88 (m, 1H, 1 × CH), 3.73-3.59 (m, 2H, 1 × $CH_2$), 3.49-3.41 (m, 1H, 1 × CH), 3.36-3.33 (m, 1H, 0.5 × $CH_2$), 3.28-3.26 (m, 1H, 0.5 × $CH_2$), 3.23 (s, 9H, 3 × $CH_3$), 2.11-1.98 (m, 2H, 1 × $CH_2$), 1.48-1.22 (m, 22H, 11 × $CH_2$), 0.90 (t, *J* = 6.8 Hz, 3H, 1 × $CH_3$). $^{13}$C NMR ($CD_3OD$, 125.77 MHz, δ): 172.5, 169.5*, 135.6, 130.9, 72.6, 67.4, 65.9, 60.5, 55.8, 54.6, 54.0, 33.6, 33.1, 30.9, 30.9, 30.9, 30.8, 30.6, 30.5, 30.5, 23.8, 14.5. MS (ESI-TOF) [m/z (%)]: 568 ([MH]$^+$, 100).

Synthesis of the membrane-forming lipids

The membrane-forming lipids (**P1** and **P2**) were chemically synthesized for NMR characterization and for verifying the corresponding chemoenzymatically generated lipids by HPLC-MS.

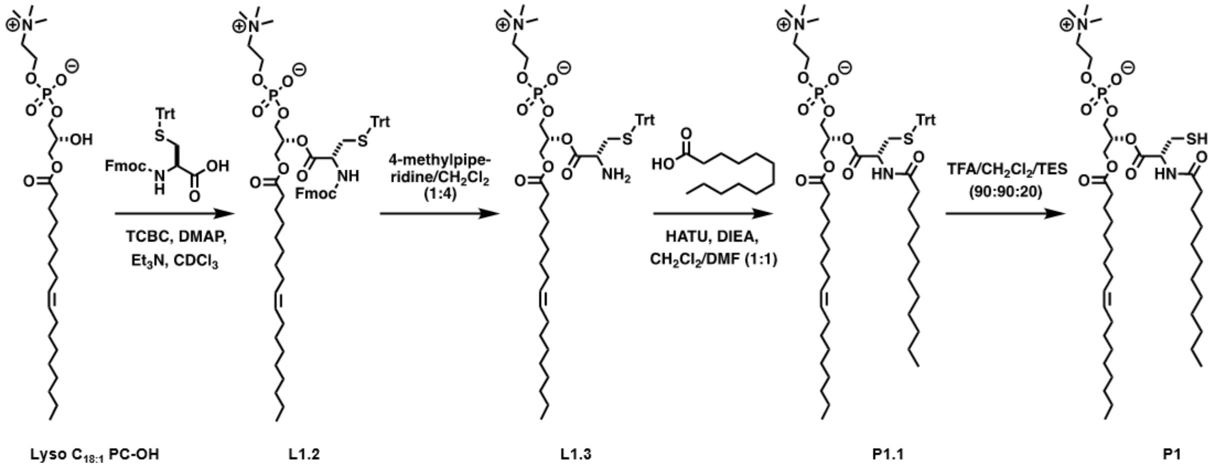


**1-oleoyl-2-[*N*-Fmoc-L-Cys(Trt)]-*sn*-glycero-3-phosphocholine (L1.2).** A solution of 1-oleoyl-2-hydroxy-*sn*-glycero-3-phosphocholine (**Lyso $C_{18:1}$ PC-OH**, 25.0 mg, 47.9 µmol), *N*-Fmoc-L-Cys(Trt)-OH (70.2 mg, 119.8 µmol), DMAP (35.1 mg, 287.5 µmol) and $Et_3N$ (23.4 µL, 167.7 µmol) in $CDCl_3$ (1.9 mL) was stirred at rt for 10 min. Then, TCBC (48.7 µL, 311.5 µmol) was added. After 12 h stirring at rt, $H_2O$ (125 µL) was added to quench the acid chloride, and the solvent was removed under reduced pressure to give a pale yellow solid. The corresponding residue was dissolved in MeOH (500 µL), filtered using a 0.2 µm syringe-driven filter, and the crude solution was purified by HPLC, affording 40.9 mg of lysophospholipid **L1.2** as a white solid [78%, $t_R$ = 8.0 min (100% *Phase B*, 25 min)]. MS (ESI-TOF) [m/z (%)]: 1111 ($[M+Na]^+$, 100), 1089 ($[MH]^+$, 72).

**1-oleoyl-2-[L-Cys(Trt)]-*sn*-glycero-3-phosphocholine (L1.3).** A solution of 1-oleoyl-2-[*N*-Fmoc-L-Cys(Trt)]-*sn*-glycero-3-phosphocholine (**L1.2**, 5.0 mg, 4.6 µmol) in 125 µL of 4-methylpiperidine/$CH_2Cl_2$ (25:100) was stirred at rt for 30 min. After removal of the solvent, the residue was dried under high vacuum for 3 h. Then, the corresponding residue was dissolved in MeOH (250 µL), filtered using a 0.2 µm syringe-driven filter, and the crude solution was purified by HPLC, affording 3.4 mg of the lysophospholipid **L1.3** as a colorless film [84%, $t_R$ = 8.4 min (50-0% *Phase A* in *Phase B*, 2.5 min, and then 100% *Phase B*, 15 min)]. MS (ESI-TOF) [m/z (%)]: 889 ($[M+Na]^+$, 60), 867 ($[MH]^+$, 100).

**1-oleoyl-2-[L-Cys(Trt)-(dodecanoyl)]-*sn*-glycero-3-phosphocholine (P1.1).** A solution of dodecanoic acid (0.5 mg, 2.3 µmol) in $CH_2Cl_2$/DMF (1:1) (200 µL) was stirred at 0 ºC for 10 min, and then HATU (1.0 mg, 2.5 µmol) and DIEA (1.6 µL, 9.2 µmol) were successively added. After 10 min stirring at 0 ºC, 1-oleoyl-2-[L-Cys(Trt)]-*sn*-glycero-3-phosphocholine (**L1.3**, 2.0 mg, 2.3 µmol) was added. After 1 h stirring at rt, the mixture was concentrated under reduced pressure. The corresponding residue was dissolved in MeOH (250 µL), filtered using a 0.2 µm syringe-driven filter, and the crude solution was purified by HPLC, affording 1.8 mg of **P1.1** as a colorless film [76%, $t_R$ = 19.7 min (50-0% *Phase A* in *Phase B*, 2.5 min, and then 100% *Phase B*, 20 min)]. MS (ESI-TOF) [m/z (%)]: 1071 ($[M+Na]^+$, 44), 1049 ($[MH]^+$, 100).

**1-oleoyl-2-[L-Cys-(dodecanoyl)]-*sn*-glycero-3-phosphocholine (P1).** A solution of 1-oleoyl-2-[L-Cys(Trt)-(dodecanoyl)]-*sn*-glycero-3-phosphocholine (**P1.1**, 1.2 mg, 1.2 µmol) in 200 µL of TFA/$CH_2Cl_2$/TES (90:90:20) was stirred at rt for 30 min. After removal of the solvent, the residue was dried under high vacuum for 3 h. Then, the corresponding residue was dissolved in MeOH (250 µL), filtered using a 0.2 µm syringe-driven filter, and the crude solution was purified by HPLC, affording 0.8 mg of the amidophospholipid **P1** as a colorless film [85%, $t_R$ = 14.4 min (50-0% *Phase A* in *Phase B*, 2.5 min, and then 100% *Phase B*, 15 min)]. $^1H$ NMR ($CDCl_3$, 500.13 MHz, δ): 6.95 (d, $J$ = 7.9 Hz, 1H, 1 × NH), 5.43-5.18 (m, 3H, 3 × CH), 4.88 (dd, $J_1$ = 8.3 Hz, $J_2$ = 4.5 Hz, 1H, 1 × CH), 4.47-4.28 (m, 3H, 1.5 × $CH_2$), 4.26-4.00 (m, 3H, 1.5 × $CH_2$),3.99-3.74 (m,

2H, 1 × $CH_2$), 3.37 (s, 9H, 3 × $CH_3$), 3.05 (ddd, $J_1$ = 13.3 Hz, $J_2$ = 8.5 Hz, $J_3$ = 4.3 Hz, 1H, 0.5 × $CH_2$), 2.91 (ddd, $J_1$ = 13.3 Hz, $J_2$ = 8.5 Hz, $J_3$ = 4.3 Hz, 1H, 0.5 × $CH_2$), 2.38-2.17 (m, 4H, 2 × $CH_2$), 2.05-1.97 (m, 4H, 2 × $CH_2$), 1.93 (br s, 1H, 1 × SH), 1.69-1.48 (m, 4H, 2 × $CH_2$), 1.37-1.16 (m, 36H, 18 × $CH_2$), 0.88 (t, $J$ = 6.8 Hz, 6H, 2 × $CH_3$). $^{13}$C NMR ($CDCl_3$, 125.77 MHz, δ): 173.7, 173.6, 169.9, 130.2, 129.9, 72.5, 66.6, 64.2, 62.5, 59.7, 54.8, 53.6, 36.5, 34.2, 32.1, 29.9, 29.9, 29.8, 29.8, 29.7, 29.7, 29.6, 29.5, 29.5, 29.4, 29.4, 29.3, 27.4, 27.3, 25.8, 25.0, 22.8, 14.3. MS (ESI-TOF) [m/z (%)]: 829 ($[M+Na]^+$, 45), 807 ($[MH]^+$, 100). HRMS (ESI-TOF) calculated for $C_{41}H_{80}N_2O_9PS$ ($[MH]^+$) 807.5317, found 807.5311.

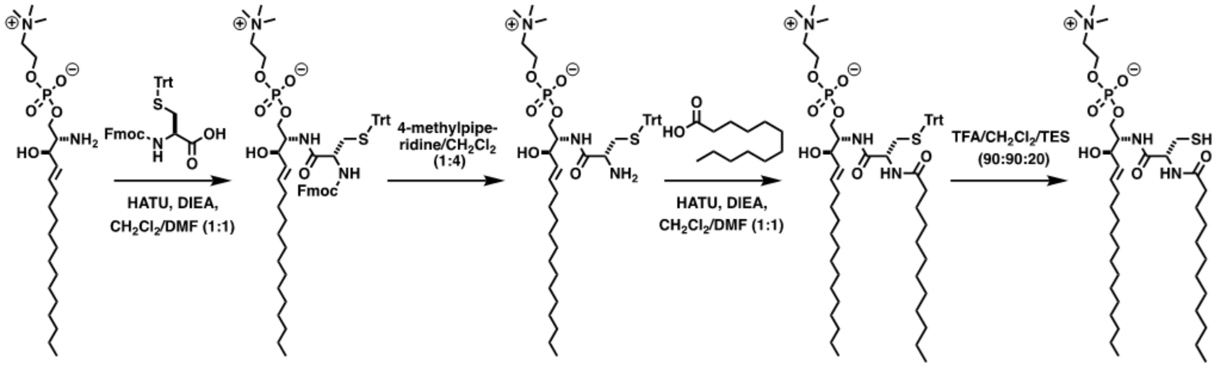


**Lyso-Sphingomyelin(d18:1)-[*N*-Fmoc-L-Cys(Trt)] (L2.2).** A solution of *N*-Fmoc-L-Cys(Trt)-OH (6.3 mg, 10.8 µmol) in $CH_2Cl_2$/DMF (1:1) (250 µL) was stirred at 0 ºC for 10 min, and then HATU (4.5 mg, 11.8 µmol) and DIEA (7.5 µL, 43.0 µmol) were successively added. After 10 min stirring at 0 ºC, lyso-sphingomyelin (d18:1) (**Lyso $d_{18:1}$ SM-$NH_2$**, 5.0 mg, 10.8 µmol) was added. After 1 h stirring at rt, the mixture was concentrated under reduced pressure. The corresponding residue was dissolved in MeOH (500 µL), filtered using a 0.2 µm syringe-driven filter, and the crude solution was purified by HPLC, affording 10.1 mg of **L2.2** as a white solid [91%, $t_R$ = 12.5 min (50-0% *Phase A* in *Phase B*, 2.5 min, and then 100% *Phase B*, 15 min)]. MS (ESI-TOF) [m/z (%)]: 1054 ($[M+Na]^+$, 23), 1032 ($[MH]^+$, 100).

**Lyso-Sphingomyelin(d18:1)-[L-Cys(Trt)] (L2.3).** A solution of Lyso-sphingomyelin (d18:1)-[*N*-Fmoc-L-Cys(Trt)] (**L2.2**, 8.0 mg, 7.8 µmol) in 125 µL of 4-methylpiperidine/$CH_2Cl_2$ (25:100) was stirred at rt for 30 min. After removal of the solvent, the residue was dried under high vacuum for 3 h. Then, the corresponding residue was dissolved in MeOH (250 µL), filtered using a 0.2 µm syringe-driven filter, and the crude solution was purified by HPLC, affording 5.3 mg of the lysosphingomyelin **L2.3** as a colorless film [84%, $t_R$ = 7.1 min (50-0% *Phase A* in *Phase B*, 2.5 min, and then 100% *Phase B*, 15 min)]. MS (ESI-TOF) [m/z (%)]: 832 ($[M+Na]^+$, 37), 810 ($[MH]^+$, 100).

**Sphingomyelin(d18:1)-[L-Cys(Trt)-(dodecanoyl)] (P2.1).** A solution of dodecanoic acid (1.0 mg, 4.9 µmol) in $CH_2Cl_2$/DMF (1:1) (250 µL) was stirred at 0 ºC for 10 min, and then HATU (2.1 mg, 5.4 µmol) and DIEA (3.4 µL, 19.8 µmol) were successively added. After 10 min stirring at 0 ºC, lyso-Sphingomyelin(d18:1)-[L-Cys(Trt)] (**L2.3**, 4.0 mg, 4.9 µmol) was added. After 1 h stirring at rt, the mixture was concentrated under reduced pressure. The corresponding residue was dissolved in MeOH (500 µL), filtered using a 0.2 µm syringe-driven filter, and the crude solution was purified by HPLC, affording 3.9 mg of **P2.1** as a colorless film [80%, $t_R$ = 15.9 min (50-0% *Phase A* in *Phase B*, 2.5 min, and then 100% *Phase B*, 15 min)]. MS (ESI-TOF) [m/z (%)]:1014 ($[M+Na]^+$, 21), 992 ($[MH]^+$, 100).

**Sphingomyelin(d18:1)-[L-Cys-(dodecanoyl)] (P2).** A solution of sphingomyelin(d18:1)-[L-Cys(Trt)-(dodecanoyl)] (**P2.1**, 2.0 mg, 2.0 µmol) in 200 µL of TFA/$CH_2Cl_2$/TES (90:90:20) was stirred at rt for 30 min. After removal of the solvent, the residue was dried under high vacuum for

3 h. Then, the corresponding residue was dissolved in MeOH (250 µL), filtered using a 0.2 µm syringe-driven filter, and the crude solution was purified by HPLC, affording 1.3 mg of the sphingomyelin **P2** as a colorless film [87%, $t_R$ = 11.8 min (50-0% *Phase A* in *Phase B*, 2.5 min, and then 100% *Phase B*, 15 min)]. $^{1}$H NMR ($CD_3OD$, 500.13 MHz, δ): 5.73 (dt, $J_1$ = 15.3 Hz, $J_2$ = 6.6 Hz, 1H, 1 × CH), 5.46 (ddt, $J_1$ = 15.3 Hz, $J_2$ = 7.6 Hz, $J_3$ = 1.5 Hz, 1H, 1 × CH), 4.51 (dd, $J_1$ = 7.7 Hz, $J_2$ = 4.9 Hz, 1H, 1 × CH), 4.36-4.20 (m, 2H, 1 × CH + 0.5 × $CH_2$), 4.18-4.04 (m, 2H, 1 × $CH_2$), 4.01-3.90 (m, 2H, 1 × CH + 0.5 × $CH_2$), 3.64 (t, $J$ = 4.7 Hz, 2H, 1 × $CH_2$), 3.22 (s, 9H, 3 × $CH_3$), 2.85 (dd, $J_1$ = 13.9 Hz, $J_2$ = 4.9 Hz, 1H, 0.5× $CH_2$), 2.74 (dd, $J_1$ = 13.8 Hz, $J_2$ = 7.7 Hz, 1H, 0.5× $CH_2$), 2.36-2.21 (m, 2H, 1 × $CH_2$), 2.10-1.97 (m, 2H, 1 × $CH_2$), 1.69-1.55 (m, 2H, 1 × $CH_2$), 1.45-1.21 (m, 38H, 19 × $CH_2$), 0.90 (t, $J$ = 6.9 Hz, 6H, 2 × $CH_3$). $^{13}$C NMR ($CD_3OD$, 125.77 MHz, δ): 174.9, 170.6, 134.0, 129.5, 70.8, 66.0, 64.1, 59.1, 55.4, 54.1, 53.2, 35.5, 32.1, 31.7, 29.5, 29.4, 29.4, 29.4, 29.4, 29.4, 29.3, 29.1, 29.1, 29.0, 28.9, 26.2, 25.6, 22.4, 13.1. MS (ESI-TOF) [m/z (%)]: 772 ($[M+Na]^+$, 32), 750 ($[MH]^+$, 100). HRMS (ESI-TOF) calculated for $C_{38}H_{77}N_3O_7PS$ ($[MH]^+$) 750.5210, found 750.5214.

#### Supplementary tables

| Fatty acid | Cys-phospholipid molecular weight | |
|---|---|---|
| | *Exact mass* | *Detected (m/z)* |
| Dodecanoic acid (C12:0) | 806.5 | 807.2 $[M+H^+]$ |
| Myristoleic acid (C14:1) | 832.5 | 833.4 $[M+H^+]$ |
| Palmitoleic acid (C16:1) | 860.6 | 861.4 $[M+H^+]$ |
| Oleic acid (C18:1) | 888.6 | 889.3 $[M+H^+]$ |

**Table S1**. Fatty acids screened for activity in this study. Formation of Cys-phospholipids was detected by HPLC-MS.

#### Supplementary figures

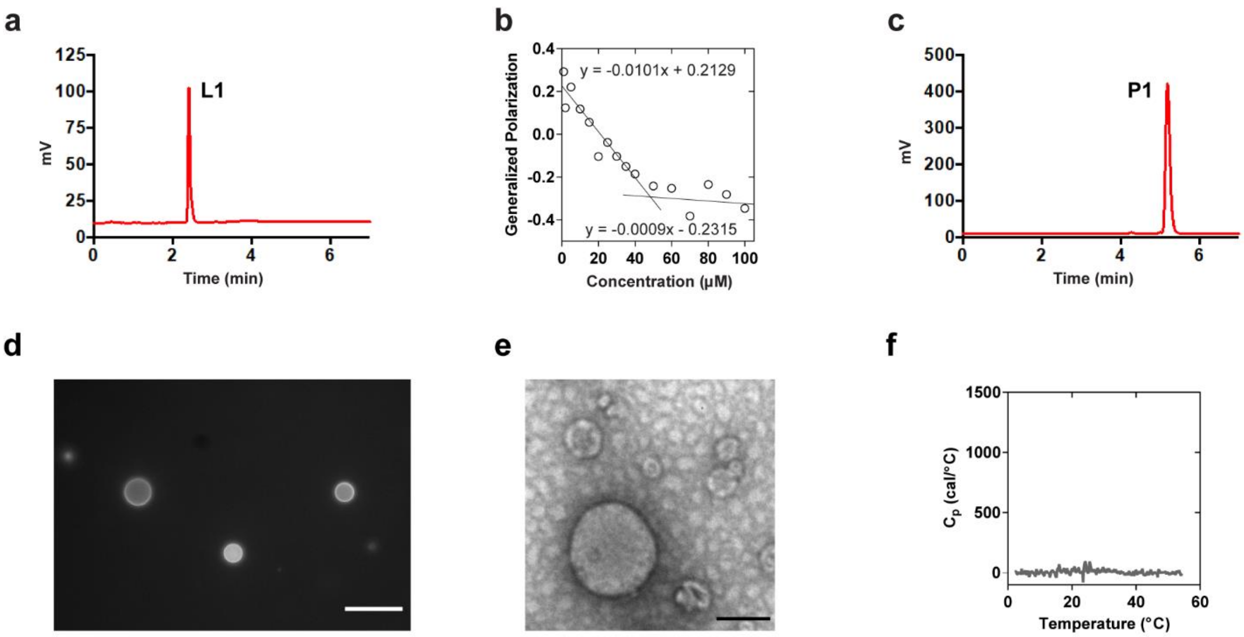


**Figure S1**. Characterization of lysolipid **L1** and phospholipid **P1**. (a) HPLC-ELSD chromatogram of pure **L1**. (b) Representative plot for critical micelle concentration (cmc) determination for lysolipid **L1** in $H_2O$ using the generalized polarization of Laurdan. The cmc was calculated to be 48 µM. (c) HPLC-ELSD chromatogram of pure **P1**. (d) Fluorescence microscopy image of vesicles of **P1** formed by gentle hydration in aqueous medium. The membranes were stained with 0.1 mol% Texas Red-DHPE. Scale bar indicates 10 µm. (e) Negative staining TEM image of vesicles formed from purified **P1** in water. Scale bar represents 100 nm. (f) Differential scanning calorimetry thermogram of a 0.25 mM aqueous dispersion of **P1** in water at scanning rate of 30 °C/h. No peaks were detected over the scan range (5-55 °C).

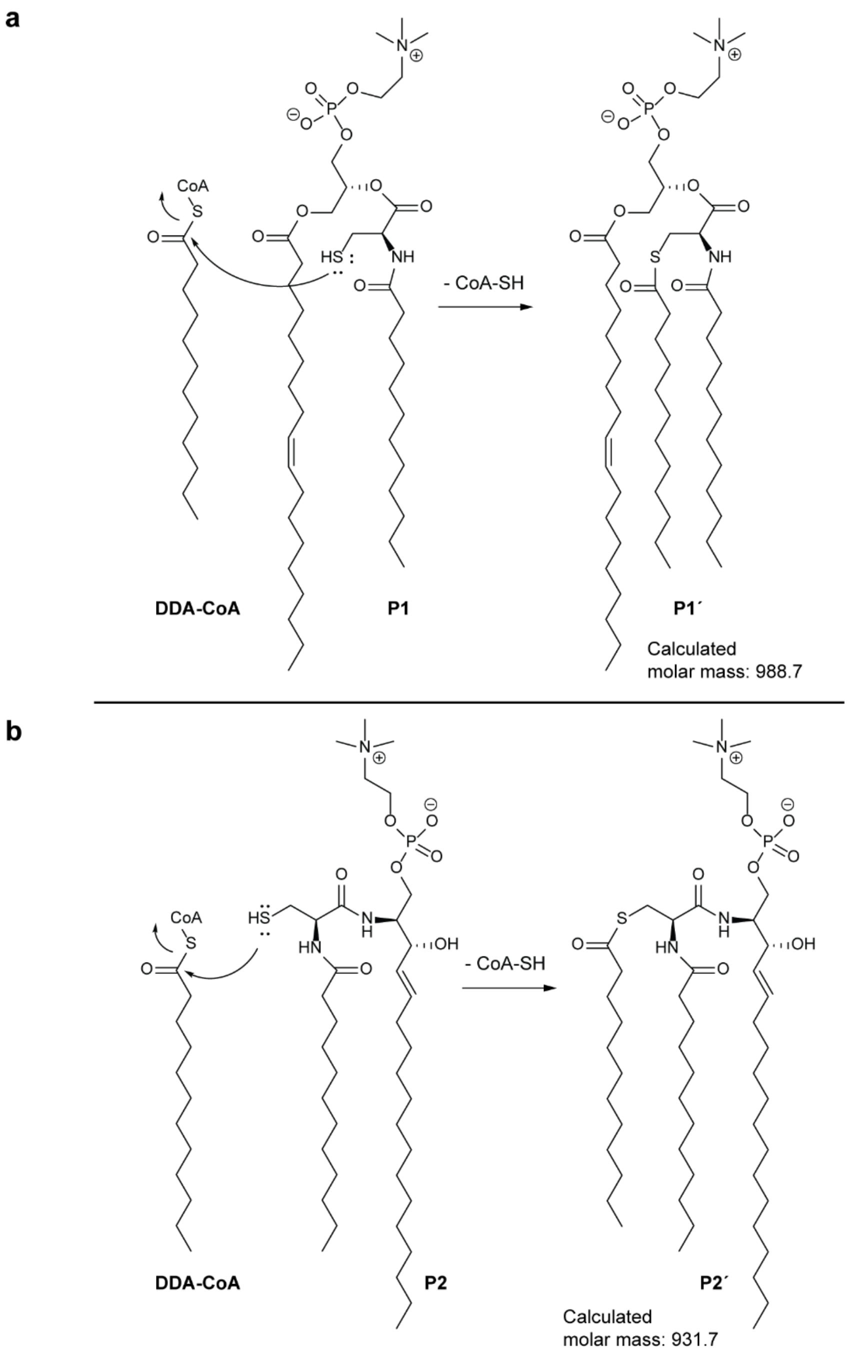


**Figure S2**. Generation of three-tailed lipids[3] by nucleophilic attack on dodecanoyl-CoA (DDA-CoA) of the free thiol (-SH) group of the two-tailed cysteine-modified (a) Phospholipid **P1** (detected m/z 514.3 corresponding to $[M+H^{+}+K^{+}]$) and (b) Sphingomyelin **P2** (detected m/z 953.8 corresponding to $[M+Na^{+}]$).

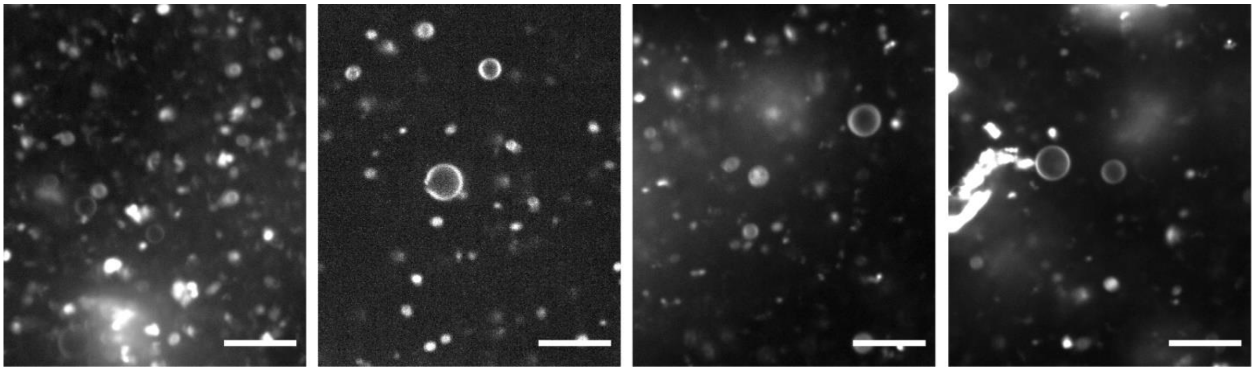

**Figure S3**. Images of FadD2-mediated in situ synthesized phospholipid **P1** vesicles in various experiments. All scale bars denote 10 µm.

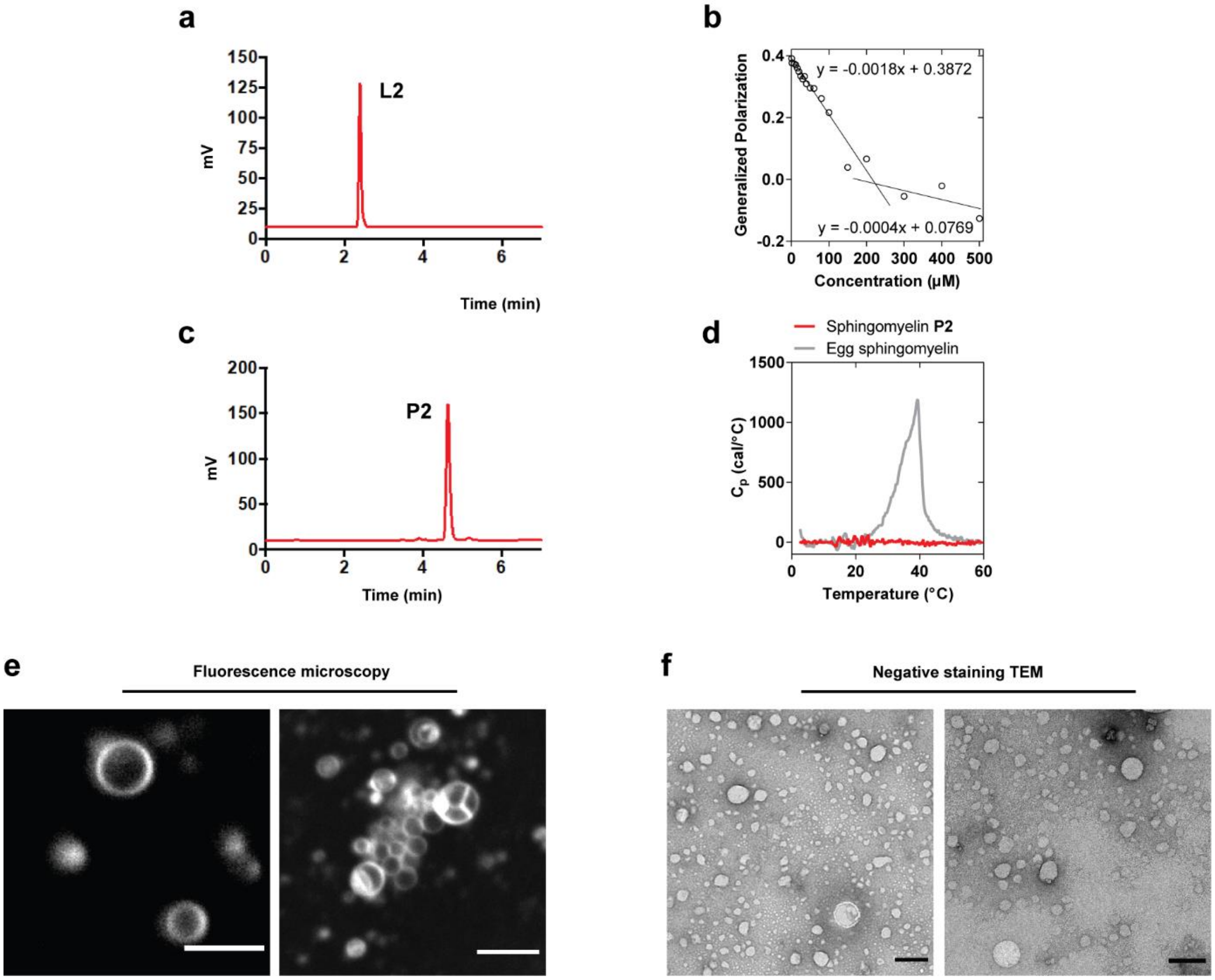


**Figure S4**. Characterization of lysolipid **L2** and sphingomyelin **P2**. (a) HPLC-ELSD chromatogram of pure **L2**. (b) Representative plot for critical micelle concentration (cmc) determination for lysolipid **L2** in HEPES-K (50 mM, pH 7.6) based on the generalized polarization of Laurdan. The cmc was calculated to be 222 µM. (c) HPLC-ELSD chromatogram of pure **P2**. (d) Differential scanning calorimetry thermogram of a 0.25 mM aqueous dispersion of **P2** in water at scanning rate of 30 °C/h. No peaks were detected over the scan range (5-60 °C). For comparison, the DSC scan of egg sphingomyelin ($T_m$ = 39.5 °C) is shown. (e) Spinning disk confocal microscopy images of vesicles obtained by gentle hydration of a film of purified **P2** in water. Membranes were stained with 0.1 mol% Nile Red. Scale bars denote 10 µm. (f) Negative staining transmission electron microscopy (TEM) images of a sonicated aqueous vesicular dispersion of pure **P2**. Scale bars denote 100 nm.

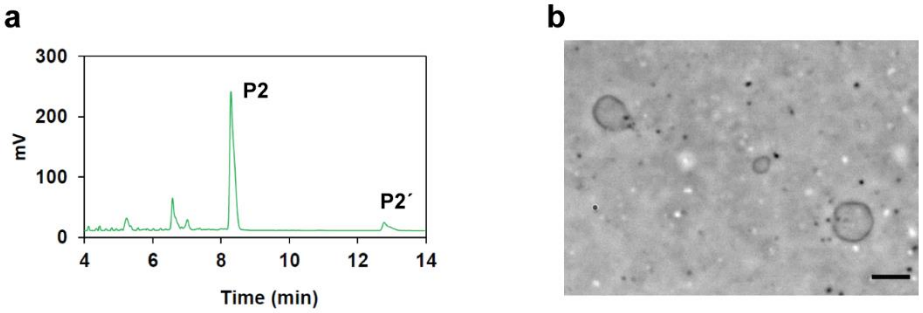


**Figure S5**. FadD2-mediated synthesis of sphingomyelin **P2** and its self-assembly into vesicles. (a) HPLC-ELSD chromatogram corresponding to synthesis of sphingomyelin **P2** mediated by FadD2. (b) Bright field microscopy of the corresponding reaction mixture showing formation of micron-sized vesicles. Scale bar represents 10 µm.

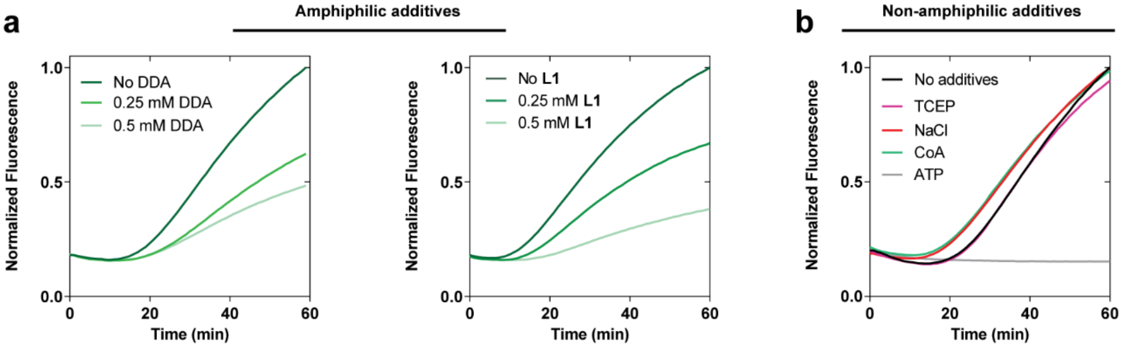


**Figure S6**. Effect of various additives on expression of sfGFP in PURE system. Effect of (a) amphiphilic additives – sodium dodecanoate (0.25 mM, 0.5 mM) and lysolipid **L1** (0.25 mM, 0.5 mM) and (b) non-amphiphilic additives TCEP (0.625 mM), NaCl (24 mM), CoA (0.64 mM), ATP (2.5 mM).

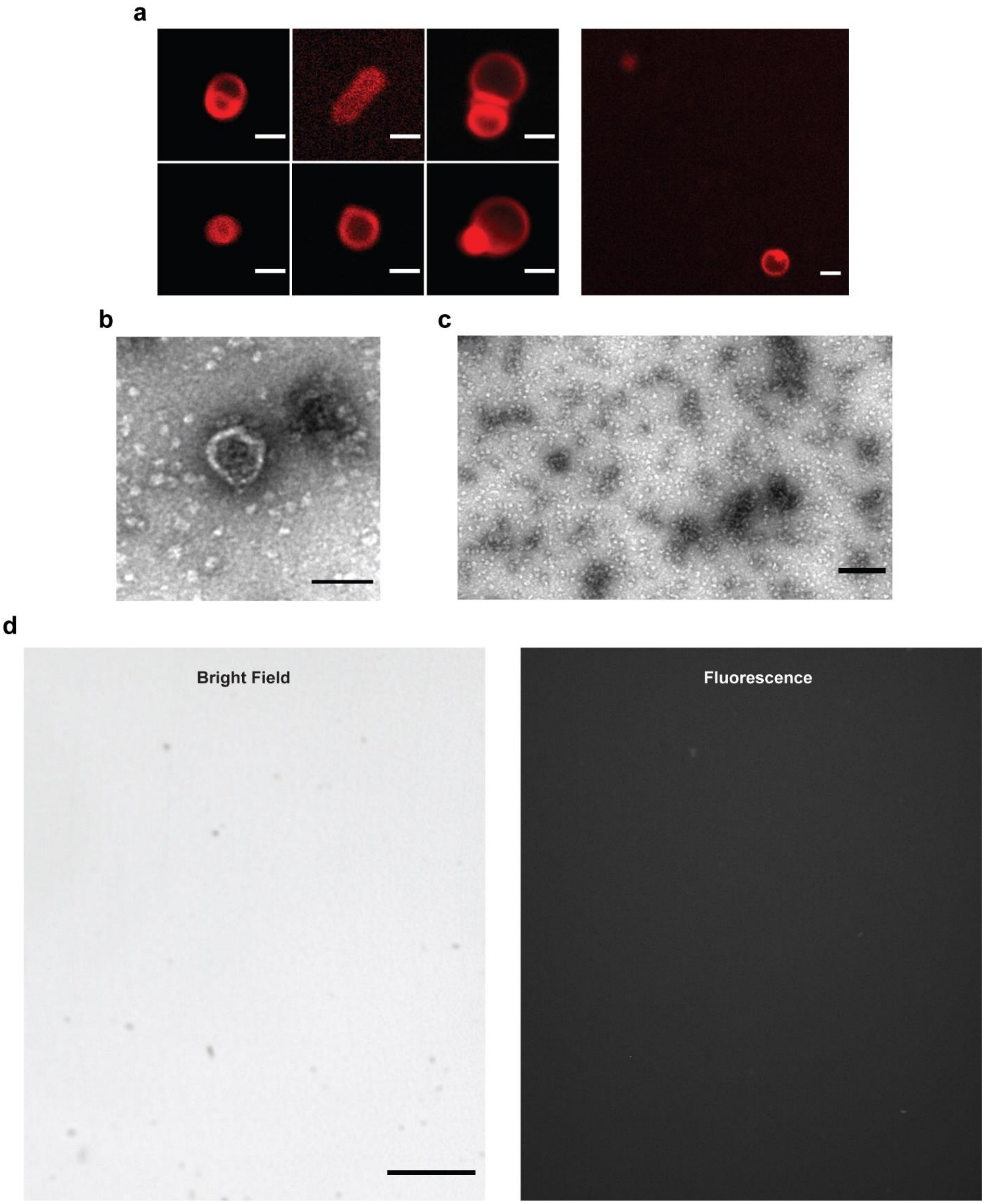


**Figure S7**. One-pot synthesis of membrane-forming lipids in PURE system. (a) Representative spinning disk confocal microscopy images of vesicles formed from sphingomyelin **P2** synthesized under one-pot condition in PURE system. All scale bars represent 2 µm. (b) Negative staining TEM of a vesicle formed from *in situ* synthesized sphingolipid **P2**. Scale bar represents 100 nm.

(c) Negative staining TEM of control condition where FadD2 DNA was omitted. Scale bar represents 200 nm. (d) Microscopy images corresponding to the control condition for one-pot reaction where the lipid precursors were incubated with no FadD2 DNA template added. The solution appeared clear with no aggregates. Scale bar denotes 20 µm.

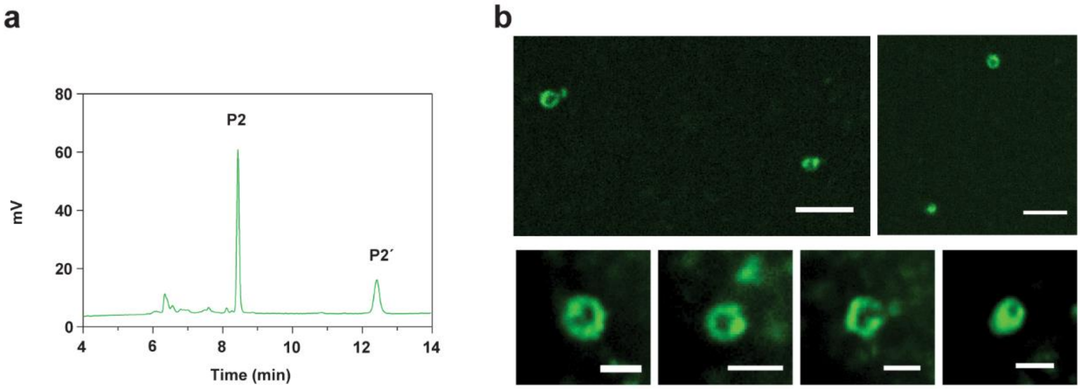


**Figure S8**. Co-expression of sfGFP-NT-lysenin and its localization to the *in situ* formed membranes. (a) HPLC-ELSD chromatogram corresponding to the one-pot reaction where FadD2 and sfGFP-NT-lysenin are co-expressed in the presence of the lipid precursors. (b) Spinning disk confocal microscopy images of *in situ* formed vesicles showing localization of sfGFP-NT-lysenin to the membranes. Scale bars in *upper* row represent 5 µm and in the *lower* row represent 2 µm.

### NMR Spectra

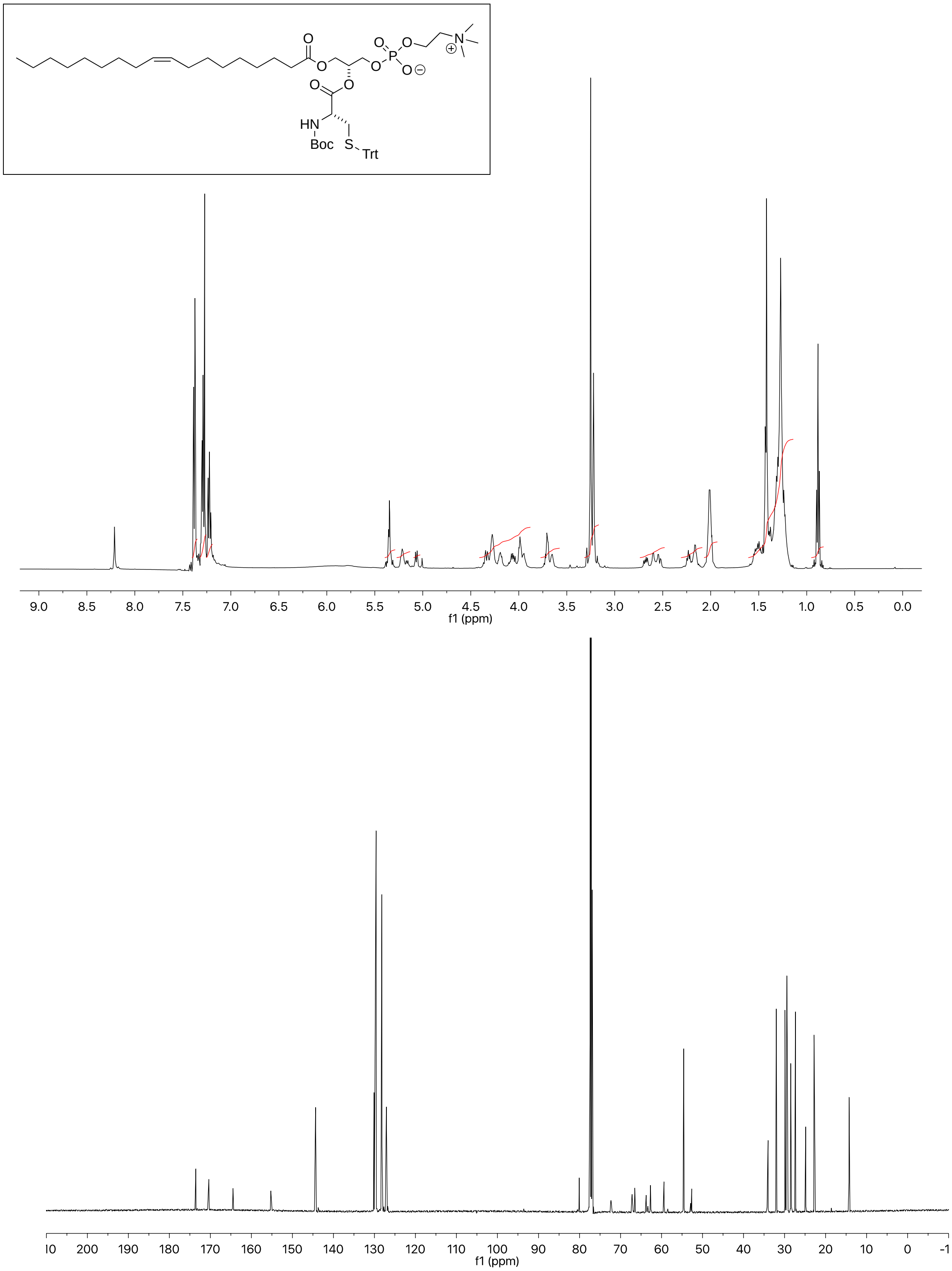


$^{1}H$ (*top*) and $^{13}C$ (*bottom*) NMR spectra of compound **L1.1**.

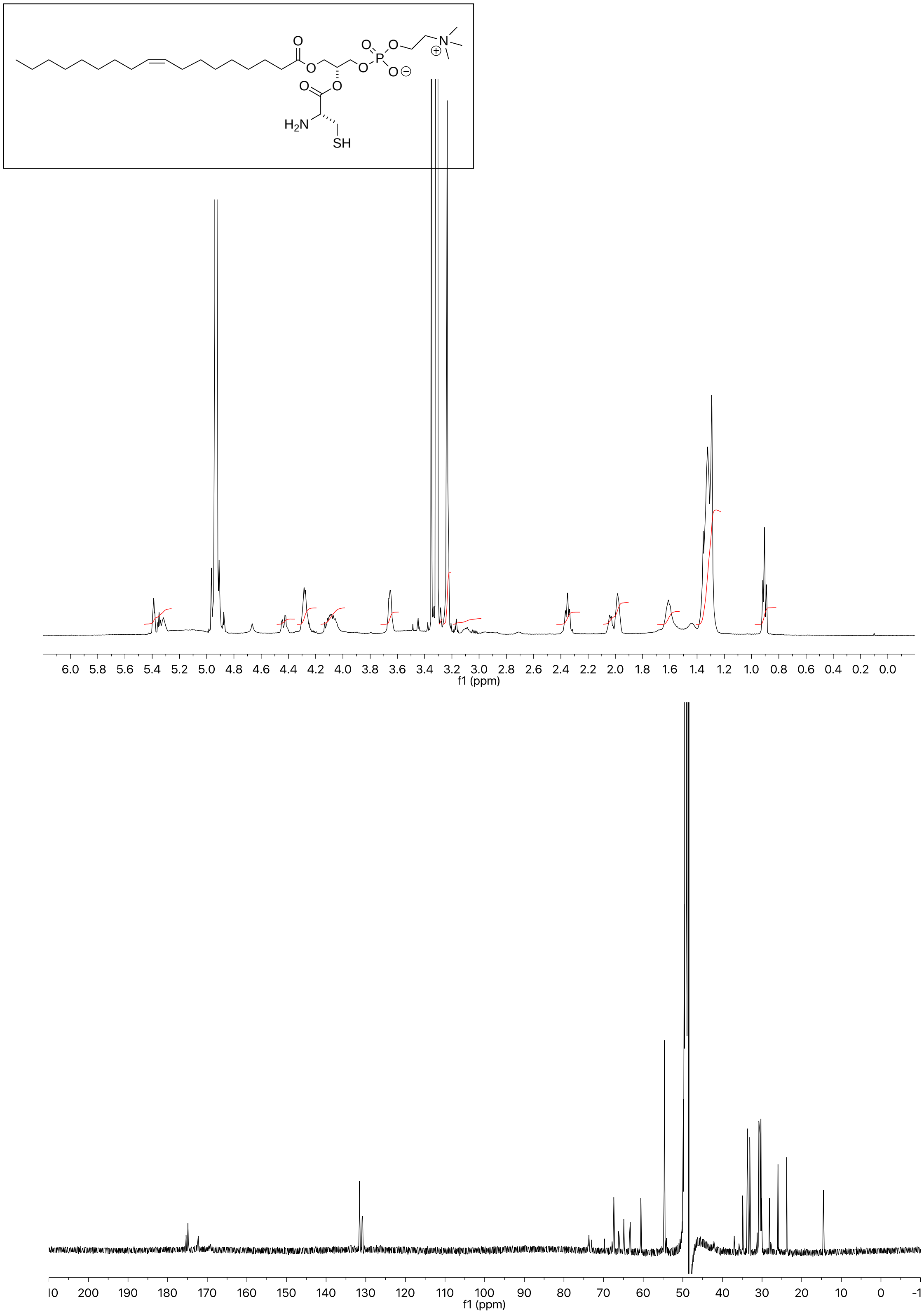


$^{1}H$ (*top*) and $^{13}C$ (*bottom*) NMR spectra of compound **L1**.

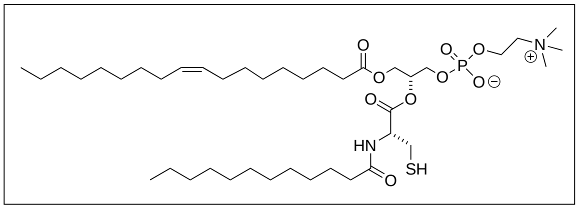


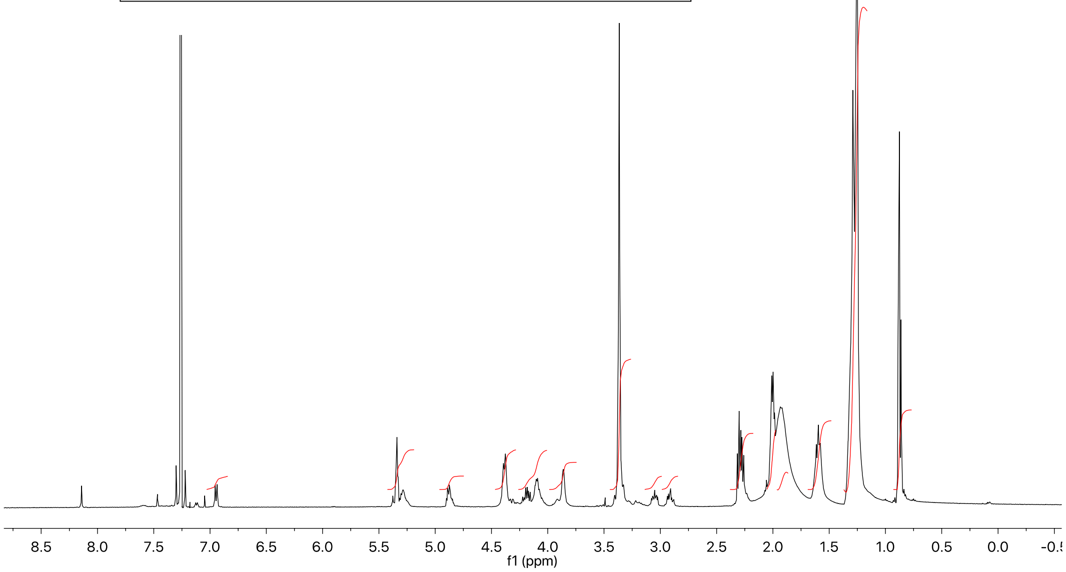


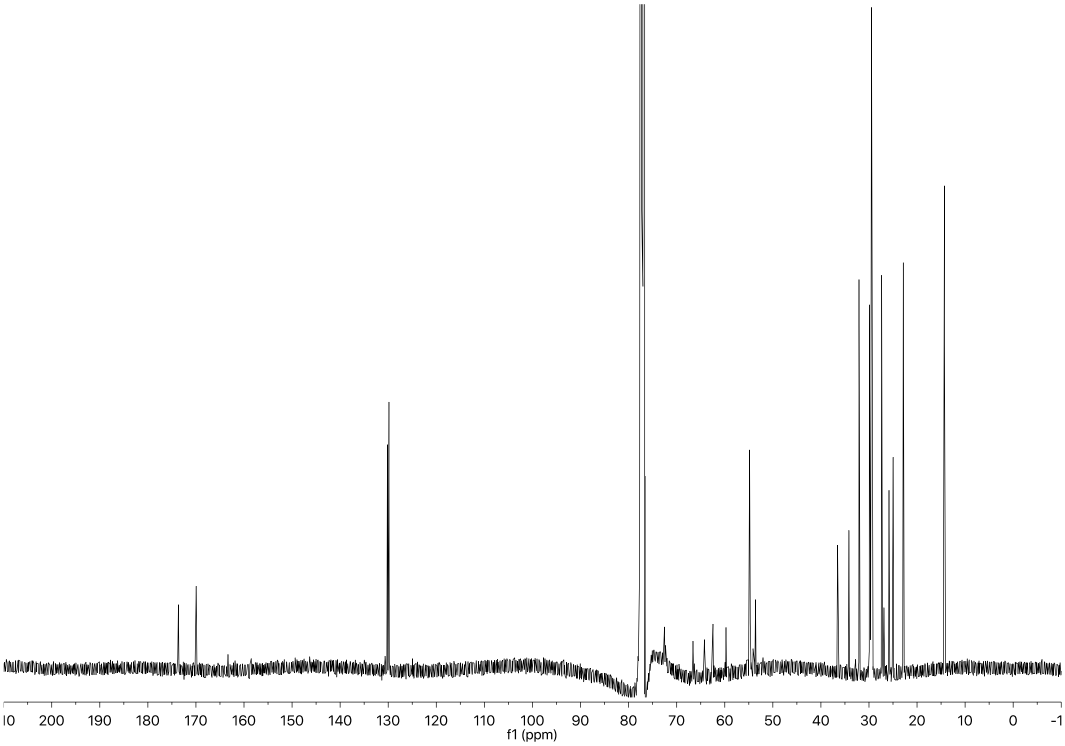


$^{1}H$ (*top*) and $^{13}C$ (*bottom*) NMR spectra of compound **P1**.

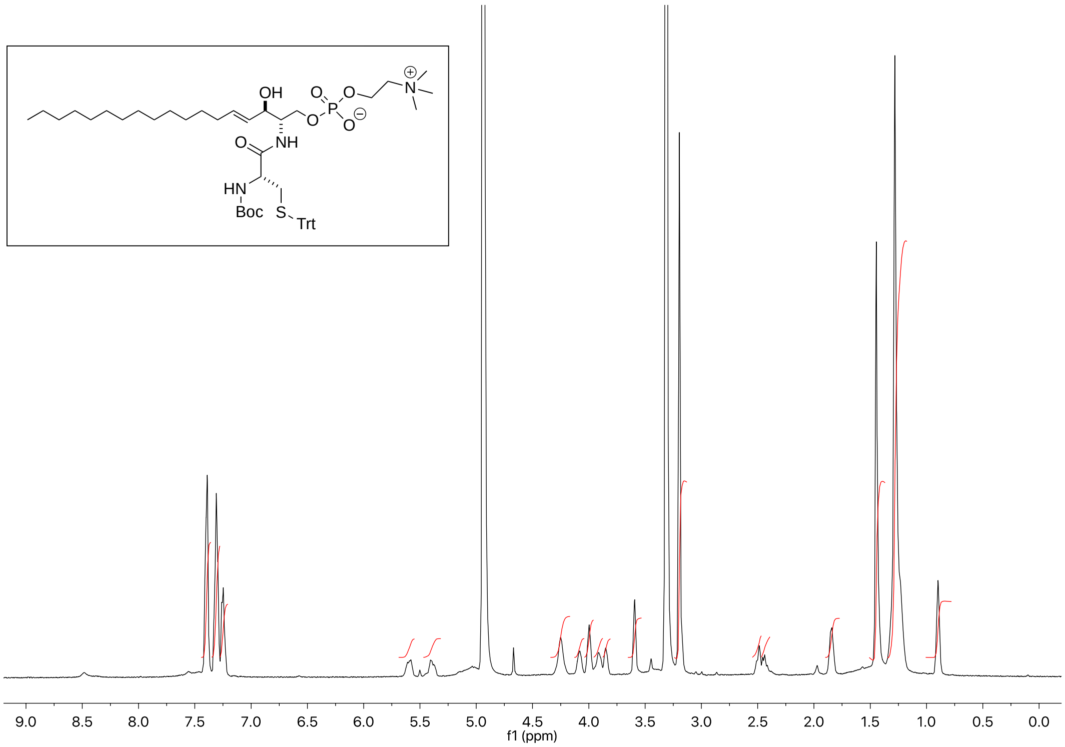


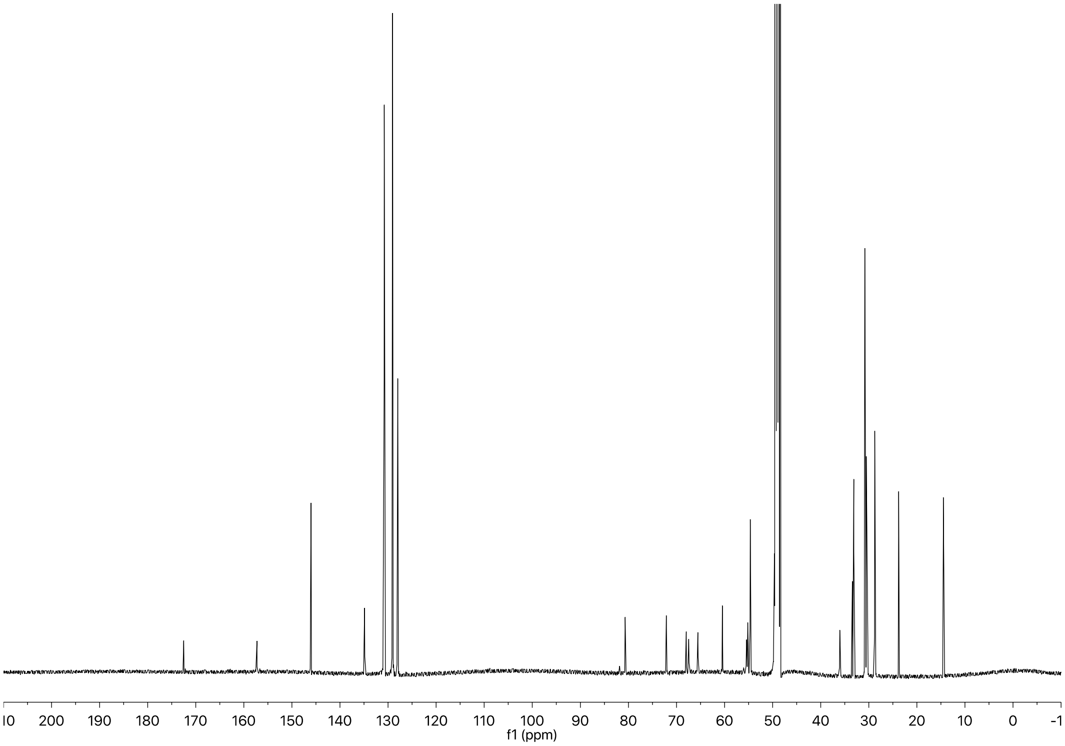


$^{1}$H (*top*) and $^{13}$C (*bottom*) NMR spectra of compound **L2.1**.

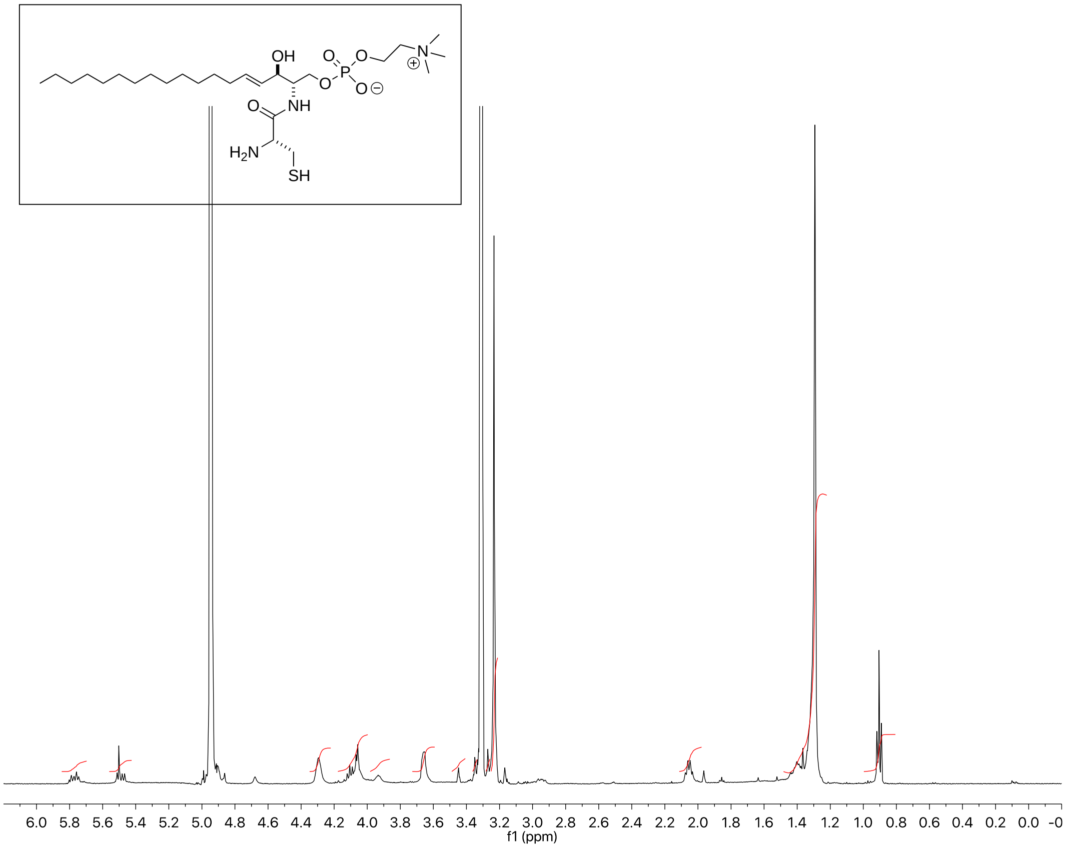


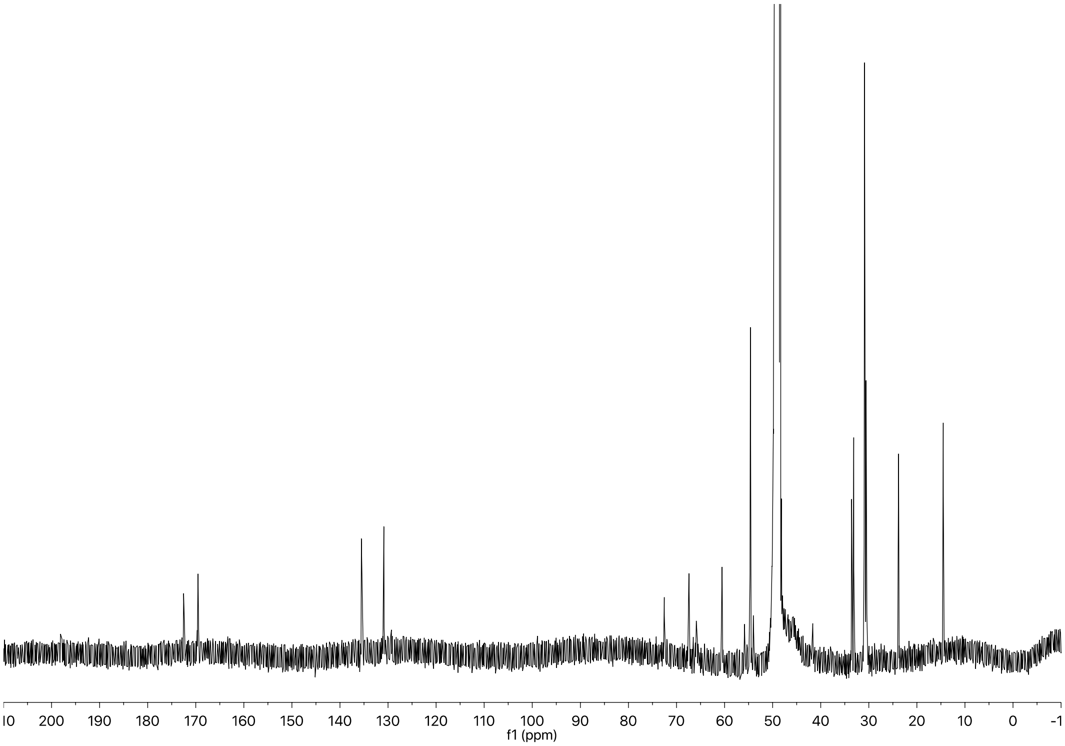


$^{1}$H (*top*) and $^{13}$C (*bottom*) NMR spectra of compound **L2**.

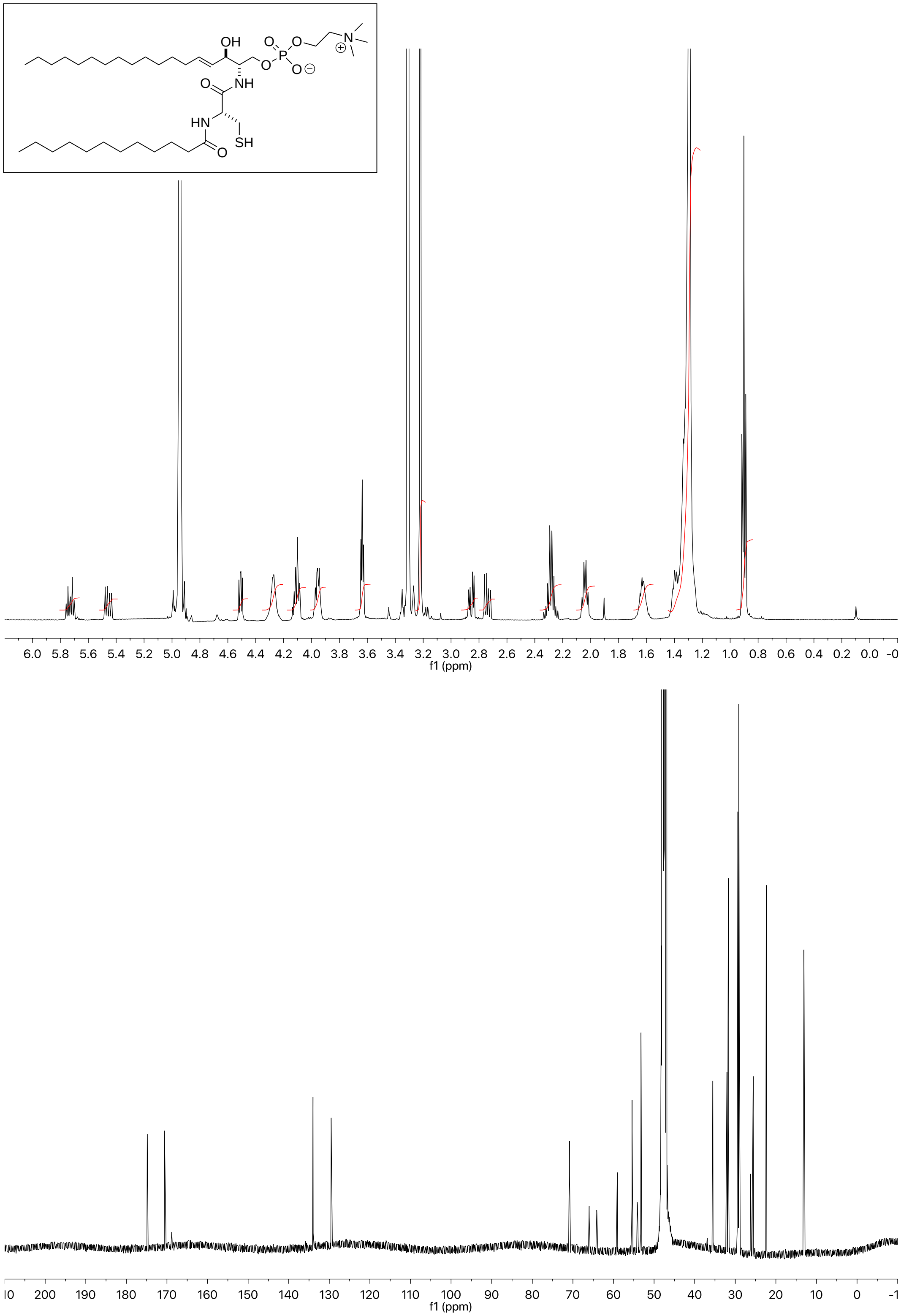


$^{1}H$ (*top*) and $^{13}C$ (*bottom*) NMR spectra of compound **P2**.
